# *L*-Arginine and asymmetric dimethylarginine (ADMA) transport across the mouse blood-brain and blood-CSF barriers: evidence of saturable transport at both interfaces and CNS to blood efflux

**DOI:** 10.1101/2024.05.30.596616

**Authors:** Mehmet Fidanboylu, Sarah Ann Thomas

## Abstract

*L*-Arginine is the physiological substrate for the nitric oxide synthase (NOS) family, which synthesises nitric oxide (NO) in endothelial and neuronal cells. NO synthesis can be inhibited by endogenous asymmetric dimethylarginine (ADMA). NO has explicit roles in cellular signalling and vasodilation. Impaired NO bioavailability represents the central feature of endothelial dysfunction associated with vascular diseases. Interestingly, dietary supplementation with *L*-arginine has been shown to alleviate endothelial dysfunctions caused by impaired NO synthesis. In this study the transport kinetics of [^3^H]-arginine and [^3^H]-ADMA into the central nervous system (CNS) were investigated using physicochemical assessment and the *in situ* brain/choroid plexus perfusion technique in anesthetized mice. Results indicated that *L*-arginine and ADMA are tripolar cationic amino acids and have a gross charge at pH 7.4 of 0.981. *L*-Arginine (0.00149±0.00016) has a lower lipophilicity than ADMA (0.00226±0.00006) as measured using octanol-saline partition coefficients. The *in situ* perfusion studies revealed that [^3^H]-arginine and [^3^H]-ADMA can cross the blood-brain barrier (BBB) and the blood-CSF barrier. [^3^H]-Arginine (11.6nM) and [^3^H]-ADMA (62.5nM) having unidirectional transfer constants (K_in_) into the frontal cortex of 5.84±0.86 and 2.49±0.35 μl.min^-1^.g^-1^, respectively, and into the CSF of 1.08±0.24 and 2.70±0.90 μl.min^-1^.g^-1^, respectively. In addition, multiple-time uptake studies revealed the presence of CNS-to-blood efflux of ADMA. Self- and cross-inhibition studies indicated the presence of transporters at the BBB and the blood-CSF barriers for both amino acids, which were shared to some degree. Importantly, these results are the first to demonstrate: (i) saturable transport of [^3^H]-ADMA at the blood-CSF barrier (choroid plexus) and (ii) a significant CNS to blood efflux of [^3^H]-ADMA. Our results suggest that the arginine paradox, in other words the clinical observation that NO-deficient patients respond well to oral supplementation with *L*-arginine even though the plasma concentration is easily sufficient to saturate endothelial NOS, could be related to ADMA transport.

## INTRODUCTION

Nitric oxide (NO) is implicated in cerebral microvessel basal tone, blood flow autoregulation and endothelium junctional integrity (1)(2). In the brain, NO participates in cell signalling and in pathogen defence, and disordered NO generation is associated with pathological conditions such as Alzheimer’s disease (3).

NO is synthesised from *L*-arginine by nitric oxide synthase (NOS) enzymes. Three isoforms of NOS have been characterised in humans (4); all of which have been classified according to the tissue or cell type of origin found when determining cDNA expression. Two of the isoforms - endothelial (eNOS) and neuronal (nNOS), have been found to be constitutively expressed, while the third isoform shown to be inducible (iNOS). Asymmetric dimethylarginine (ADMA) is a biologically significant inhibitor of NO production as it cannot be used as a substrate by NOS (5,6).

As *L*-arginine is the precursor for NO synthesis (7) and ADMA a potent endogenous inhibitor of NO synthesis (5) (6), the interplay between the transport of these two cationic amino acids at the blood-brain barrier (BBB) and blood-cerebrospinal fluid (CSF) barriers is likely to directly relate to NO production at these interfaces and within the central nervous system (CNS). In fact, the arginine/ADMA plasma concentration ratio is widely considered to be an important indicator of NO bioavailability. Interestingly, dietary supplementation with *L*-arginine has been shown to alleviate endothelial dysfunctions caused by impaired NO synthesis (8)(9), despite eNOS already being saturated with *L*-arginine (10)(11). This is called the arginine paradox. Taking into consideration that ADMA can enter and exit cells (12) (13), understanding ADMA and *L*-arginine transport in more detail at the blood-brain and blood-CSF barriers will contribute to further understanding the pathway for NO generation and how it is potentially altered by disease and supplements. We have already investigated [^3^H]-arginine and [^3^H]-ADMA transport using an *in vitro* model of the BBB (hCMEC/D3 cells) (14). In this present study we (i) utilized a chemical property database to compare the physicochemical characteristics of the cationic amino acids and (ii) examined the transport of [^3^H]-arginine and [^3^H]-ADMA using an *in situ* brain /choroid plexus perfusion method in anaesthetized mice. This sensitive method allows the transport processes at the BBB (so the brain capillary wall) to be distinguished from those at the blood-CSF interface (so the choroid plexuses and the arachnoid membrane). We will also explore the hypothesis that ADMA efflux could be responsible for the increase in NO production after arginine supplementation by performing self- and cross-inhibitor studies.

## METHODS

### Materials

[^3^H]-arginine (mol. wt., 174.2 g/mol; specific activity, 43 Ci/mmol; >97% radiochemical purity) was purchased from Amersham Radiochemicals, Buckinghamshire, UK. [^3^H]-ADMA (mol. wt., 276.7 g/mol; specific activity, 8 Ci/mmol; 96.4% radiochemical purity) was synthesized and tritiated by Amersham Radiochemicals, Cardiff, UK. [^14^C]-sucrose (mol. wt., 342.3 g/mol; specific activity, 0.412 Ci/mmol; 99% radiochemical purity; Moravek Biochemicals, Brea, CA). Unlabelled *L*-arginine and N^G^,N^G^-dimethylarginine dihydrochloride (ADMA) were purchased from Sigma Aldrich (Dorset, UK).

### Physicochemical characteristics

The physicochemical properties of *L*-arginine, ADMA and sucrose were obtained from the chemical properties database, MarvinSketch (15). These properties included their chemical structures, the percentage distribution of the different microspecies at physiological pH, molecular weight, predicted log D at pH 7.4 and the gross charge distribution at pH 7.4.

The lipophilicity of the radiolabelled test molecules was also measured by means of an octanol:saline partition coefficient. 0.75 mL phosphate buffered saline (pH 7.4) containing 1 μCi (0.037 MBq) of either [^14^C]-sucrose (2.5 μM), [^3^H]-arginine (101.0 nM) or [^3^H]-ADMA (542.8 nM) was mixed with 0.75 mL octanol (Sigma Aldrich; Dorset, UK). Centrifugation separated the hydrophobic (octanol) phase and hydrophilic (saline) phase and 100 μL samples in triplicate were taken for radioactive liquid scintillation counting on a Tricarb 2900TR liquid scintillation counter (Perkin-Elmer; Boston, MA, USA). The partition coefficient was calculated as the ratio of radioactivity in the octanol phase to the radioactivity in the aqueous phase. The values for *L*-arginine and ADMA were compared.

### Animals & anaesthesia

All experiments requiring the use of animals were performed in accordance with the Animal (Scientific Procedures) Act, 1986 and Amendment Regulations 2012 and with consideration to the Animal Research: Reporting of *In Vivo* Experiments (ARRIVE) guidelines. The study was approved by the King’s College London Animal Welfare and Ethical Review Body.

All animals used in procedures were adult male BALB/c mice (between 23 g and 25 g) sourced from Harlan Laboratories, Oxon, UK, unless otherwise stated. Free access to water and food was provided *ad libitum*. Domitor® (medetomidine hydrochloride) and Vetalar® (ketamine) were both purchased from Harlan Laboratories; Cambridge, UK. All animals were anaesthetised (2 mg/kg Domitor® and 150 mg/kg Vetalar® injected intraperitoneally), and a lack of self-righting and paw-withdrawal reflexes (as surrogate indicators of consciousness) thoroughly checked prior to carrying out all procedures. Animals were heparinised with 100 units heparin (in 0.9% m/v NaCl_(aq)_, Harlan Laboratories; Oxon, UK), administered *via* the intraperitoneal route prior to surgery.

### *In situ* brain/choroid plexus perfusion

The *in situ* brain/choroid plexus perfusion method is designed to directly perfuse the brain with artificial plasma *via* a cannula placed in the heart. Artificial plasma is prepared to mimic the ionic composition of blood, and transporter inhibitors can be added in known concentrations and delivered to the brain for defined periods of time. There are a number of advantages associated with using the *in situ* brain/choroid plexus perfusion technique over other methods (e.g. brain uptake index method) to study blood-brain and blood-CSF barrier transport. These advantages include:

**i.** Perfusions can be carried out over a long period of time (up to 30 minutes), which allows measurement of the CNS uptake of slowly permeating molecules.
**ii.** Heart perfusions allow transport across the BBB and blood-CSF barrier to be measured simultaneously.
**iii.** The solute of interest is delivered to the CNS at a known concentration with a constant flow that does not vary with time and the kinetics of transport can be calculated accordingly.
**iv.** The concentration of transporter inhibitors in the artificial plasma can easily be manipulated to measure saturable transport across the BBB and blood-CSF barrier and identify the transport systems involved.
**v.** The solute is exposed to the brain microcirculation before any peripheral organs involved in metabolism (*e.g.* liver, kidneys or lungs) and thus one can be confident that the compound is intact when interacting with transporters at the blood-brain and blood-CSF barriers.

#### Surgical preparation

The experimental procedure followed a previously outlined protocol (16). The left ventricle of the heart was cannulated with a 25G x 10 mm butterfly-winged needle connected to a perfusion circuit. An artificial plasma was warmed to 37°C and oxygenated (95% O_2_ and 5% CO_2_) before being perfused into the heart *via* this circuit at a flow rate of 5 mL/min. The right atrium of the heart was sectioned to allow outflow of the artificial plasma. Thus, an open circuit is created and the artificial plasma only passes through the circulation once. The artificial plasma consisted of a modified Krebs-Henseleit mammalian Ringer solution with the following constituents: 117 mM NaCl, 4.7 mM KCl, 2.5 mM CaCl_2_, 1.2 mM MgSO_4_, 24.8 mM NaHCO_3_, 1.2 mM KH_2_PO_4_, 10 mM glucose, and 1 g/liter bovine serum albumin. Radiolabelled and unlabelled test molecules were infused into the inflowing artificial plasma for periods up to 30 minutes.

#### CNS sampling

At the set perfusion time a cisterna magna CSF sample was taken, the animal decapitated and the brain removed. Specific brain samples including the circumventricular organs (CVOs) were taken using a Leica S4E L2 stereomicroscope (Leica; Buckinghamshire, UK). The brain samples included the frontal cortex, caudate nucleus, occipital cortex, hippocampus, hypothalamus, thalamus, pons and cerebellum. The CVOs included the choroid plexus, pituitary gland and pineal gland.

#### Capillary depletion analysis

The microvasculature within a sample of brain tissue can be separated from the brain parenchyma using capillary depletion analysis (17). Following perfusion, the remaining brain tissue after microdissection of the regions described above (typically 200-300 mg) underwent capillary depletion following a method modified for mouse (16). A whole-brain homogenate was prepared using physiological capillary depletion buffer (141 mM NaCl, 4.0 mM KCl, 2.8 mM CaCl_2_, 1.0 mM MgSO_4_, 10.9 mM HEPES, 1.0 mM NaH_2_PO_4_, 10 mM glucose at pH 7.4) and a manual homogeniser, before adding 26% m/v dextran (final concentration 13% m/v, 60-90kDa, MP Biomedicals Europe, France) and homogenising again. Both the physiological capillary depletion buffer and dextran solution were maintained at 4 °C both to halt any cell metabolism and transport processes, and to reduce any cellular or protein damage due to heat generated from the homogenisation process. The homogenate was centrifuged at 5,400 r.c.f and 4°C to produce an endothelial cell-enriched pellet and supernatant containing brain parenchyma.

#### Liquid scintillation analysis

All samples (brain regions, capillary depletion brain homogenate, pellet and supernatant, CVOs, CSF and plasma samples) were solubilised in 0.5 mL tissue solubiliser (Solvable; Perkin-Elmer; Boston, MA, USA). After incubating the samples at room temperature for 48 h, 4 mL scintillation fluid (Lumasafe®; Perkin-Elmer; Boston, MA, USA) was added to each before vigorous vortexing. The amounts of [^3^H] and [^14^C] radioactivity in each sample were then quantified using a Packard Tri-Carb 2900TR counter (Perkin-Elmer; Boston, MA, USA). Counts per minute were then converted to disintegrations per minute (dpm) by the counter using internally stored quench curves from standards and corrected for background dpm.

### Experiment design

The *in situ* brain perfusion experiments contained [^3^H]-arginine (11.6 nM) or [^3^H]-ADMA (62.5 nM) and [^14^C]sucrose in the artificial plasma. Multiple-time (2.5, 10, 20 and 30 minute) uptake studies were performed to measure the uptake of [^3^H]-arginine or [^3^H]-ADMA into the mouse CNS over time. Self-inhibition experiments were also performed. In these experiments the perfusion period was 10 minutes and [^3^H]-arginine was measured in the presence of 100 μM unlabelled *L*-arginine and the uptake of [^3^H]-ADMA measured in the presence of 100 μM unlabelled ADMA. Cross-competition experiments were also performed. In these experiments the perfusion period was 10 minutes and [^3^H]-arginine was measured in the presence of 0.5-500 μM unlabelled ADMA and the uptake of [^3^H]-ADMA measured in the presence of 100 μM arginine.

[^14^C]-Sucrose is used as an internal control and determines blood-brain and blood-CSF barrier integrity in each experiment. It can also provide a measure of the cerebrovascular space in different regions, and the extracellular space formed between choroid plexus capillary endothelium and epithelium (18). In the pituitary gland and pineal gland samples, [^14^C]-sucrose represents the vascular space and the ability of [^14^C]-sucrose to cross between capillary endothelial cells in the absence of tight junctions.

Values for uptake of [^3^H]-arginine or [^3^H]-ADMA in brain regions can be corrected for [^14^C]-sucrose vascular space to standardize uptake and reduce regional discrepancies due to differences in vascularity. It also corrects for individual variation. Thus, any differences in the brain uptake of radiolabelled solutes are likely to be due to differences in transporter expression, rather than due to differences in blood supply.

### Expression of results

The concentration of radioactivity in the brain regions (C_BRAIN_ ; dpm/g), CVOs (C_CVO_ ; dpm/g) and CSF (C_CSF_ ; dpm/mL) was expressed as a percentage of that in the artificial plasma (C_pl_; dpm/mL) and termed *R_BRAIN_*(dpm/100g), *R_CVO_* (dpm/100g) or *R_CSF_* (dpm/100mL), respectively, or R_Tissue_ (dpm/100g) as shown in equation 1:

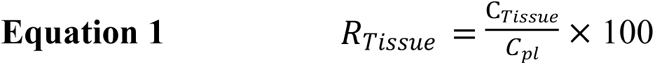

Correcting for vascular space involved subtracting the *R_Tissue_* value obtained for [^14^C]-sucrose in each sample from the *R_Tissue_* value concurrently obtained for the [^3^H]-labelled cationic amino acid.

Using these data, the permeability of the BBB and blood-CSF barrier to the solute of interest can be calculated as a unidirectional transfer constant (*K_in_*). There are two methods to determine the K*_in_* value:

i. Single-time point analysis method, described by equation 2
ii. Multiple-time point analysis method, described by equation 3

The first method: (i) was developed by (19,20) and applied to the brain perfusion data (21).

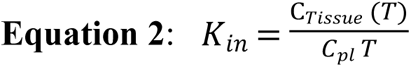

Where C_Tissue_ (T) is radioactivity (dpm) per g of tissue at time-point T (perfusion time in minutes), and C_pl_ is radioactivity (dpm) per mL of artificial plasma.

Equation 2 assumes that the entry of the radiolabelled solute of interest into the CNS is proportional to, but less than, its concentration in the artificial plasma and efflux (CNS to blood) is much smaller than influx (blood to CNS) of the test solute and therefore can be ignored (22).

It should however be noted that calculating the K_in_ value from blood to CNS using this method requires that *R_Tissue_* at time T is first corrected for vascular space by subtracting the *R_Tissue_* value for [^14^C]-sucrose determined at that time-point.

The second method (ii) for calculating the K_in_ value from blood to CNS requires multiple-time uptake data and equation 3 (23–25)(21):

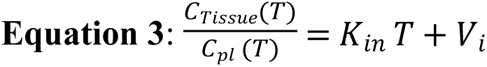

Where C_Tissue_ (T) and C_pl_ (T) are radioactivities per unit weight of tissue and plasma at time T; T being the length of perfusion. *V_i_* is the initial volume of distribution of the test solute in the rapidly equilibrating space (which may include the vascular space, the capillary endothelial volume and /or compartments in parallel with the BBB, such as the space between the brain capillary endothelium and the neuroglial lining of the capillary) (21,22,26). Thus, plotting the data points resulting from **equation 1** at each time point defines a straight line (*y=mx+c*), where *K_in_* is the slope, and *V_i_* is the ordinate intercept. Any transport of the radiolabelled solute of interest back from the brain to the blood can be detected by a loss of linearity of the experimental points (21). During the experimental period when equation 3 is applicable, the amount of test substance in V*_i_* is roughly proportional to C_pl_ and the test substance moves unidirectionally from plasma into brain tissue (26).

The percentage change in the R_Tissue_ uptake values achieved in the absence or presence of an unlabelled inhibitor can be determined by means of equation 4.

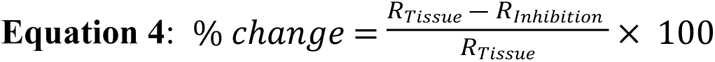

We define R_Inhibition_ as the R_Tissue_ uptake in the presence of an inhibitor in the artificial plasma. A decrease in the uptake of radiolabelled solute in the presence of unlabelled inhibitor is indicative of a saturable influx transport system. Conversely, an increase in the uptake of radiolabelled solute in the presence of unlabelled inhibitor is indicative of a saturable efflux transport system. No change in the distribution of the radiolabelled solute in the absence or presence of the unlabelled inhibitor may indicate the absence of saturable transport by the radiolabelled solute or the use of influx and efflux transporters by the radiolabelled solute.

### Statistics

Data from all experiments are presented as mean ± standard error of the mean (SEM). Statistical significance was taken as follows: not significant (ns), *p* > 0.05, \**p* < 0.05, \*\**p* < 0.01, \*\*\**p* < 0.001. All statistical analyses were performed using GraphPad Prism v5.0c or v6 graphing and statistics package for Mac.

## RESULTS

### Physicochemical characteristics

The physicochemical characteristics of *L*-arginine and ADMA were obtained from the chemical properties database, MarvinSketch (15). *L-*Arginine and ADMA are cationic (tripolar) amino acids, which exist as two microspecies at physiological pH. The percentage distribution of the two microspecies for each amino acid is shown in Fig 1. The major microspecies of both *L*-arginine (98.15%) and ADMA (98.11%) has a positive charge (+1). The minor microspecies of both *L*-arginine (1.85%) and ADMA (1.88%) are zwitterions. Both *L*-arginine and ADMA have a gross charge at physiological pH of +0.981. The molecular weights of arginine and ADMA are 174.2 g/mol and 202.26 g/mol respectively. The log D at pH 7.4 of arginine is −4.77 and ADMA is −3.99 (15). Octanol-saline partition coefficients at pH 7.4 of [^3^H]-arginine and [^3^H]-ADMA were also measured as part of this study and were found to be 0.00149±0.00016 and 0.00226±0.00006, respectively.

**Fig 1:**
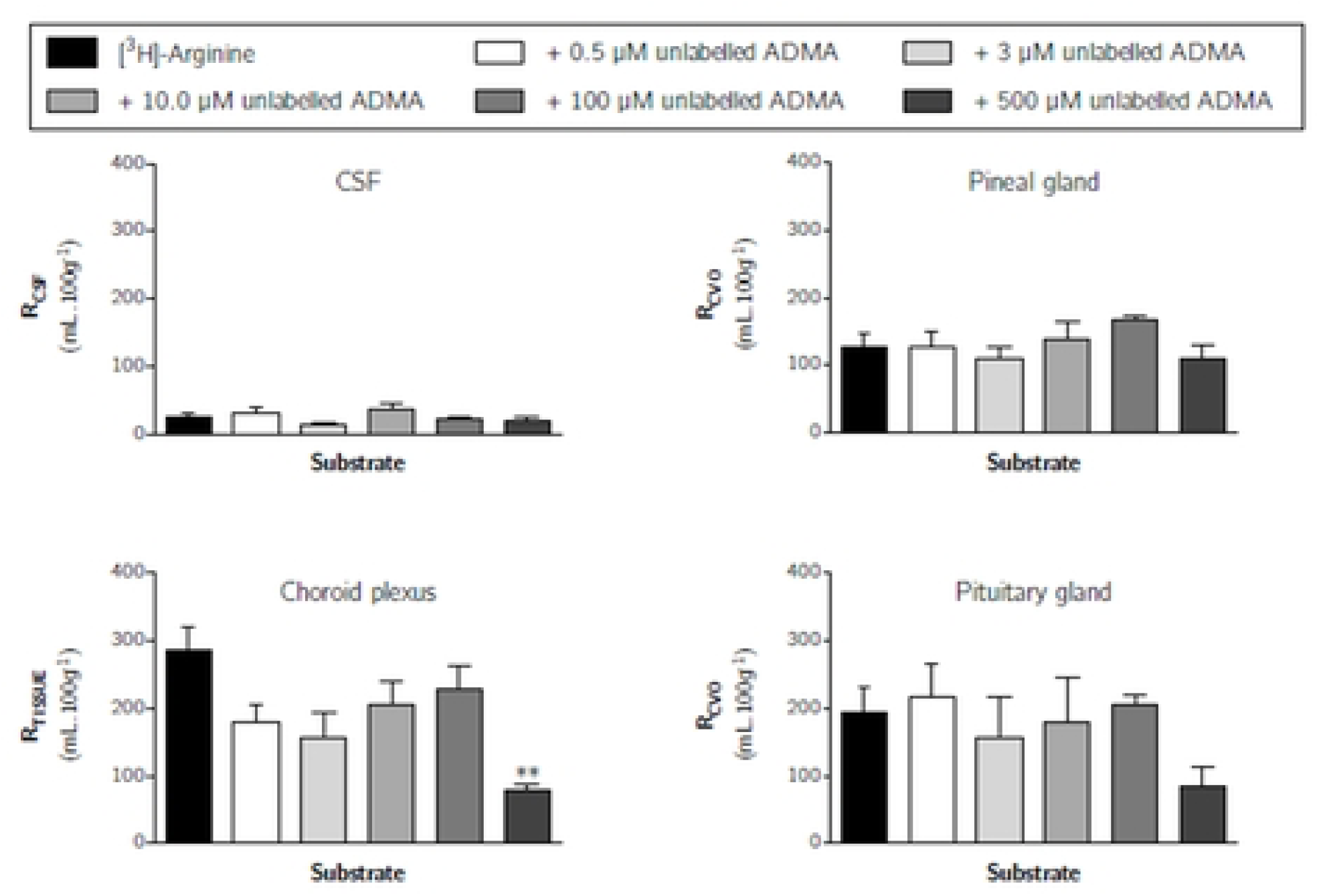
The percentage distribution and chemical structures of the two *L*-arginine microspecies and the two ADMA microspecies found at physiological pH. Microspecies A is the major microspecies. Microspecies B is the minor microspecies.

[^14^C]-Sucrose is used as a baseline marker molecule. It has a molecular weight of 342.3 g/mol and it has a log D at pH 7.4 of −4.87 (15). The octanol-saline partition coefficient at pH 7.4 of [^14^C]-sucrose was 0.00105±0.00022.

### Arginine

#### Brain distribution

Fig 2 shows the distribution (R_BRAIN_) of the vascular space marker molecule, [^14^C]-sucrose, and the cationic amino acid, [^3^H]-arginine, into different regions of the brain over time as measured by the *in situ* brain/choroid plexus perfusion method. Multiple time uptake analysis of [^14^C]-sucrose into the different brain regions was used to calculate a unidirectional transfer constant (K*_in_*) and an initial volume of distribution (V*_i_*) by means of equation 3 and the values are reported in Table 1. For example, the K*_in_* was 0.070±0.016 μl.min^-1^.g^-1^ and the V*_i_* was 1.40±0.31 mL.100g^-1^ for [^14^C]-sucrose distribution into the frontal cortex. The low distribution of [^14^C]-sucrose into all regions of the brain and at all time points after *in situ* perfusion confirms the integrity of the BBB in each experiment.

**Fig 2:**
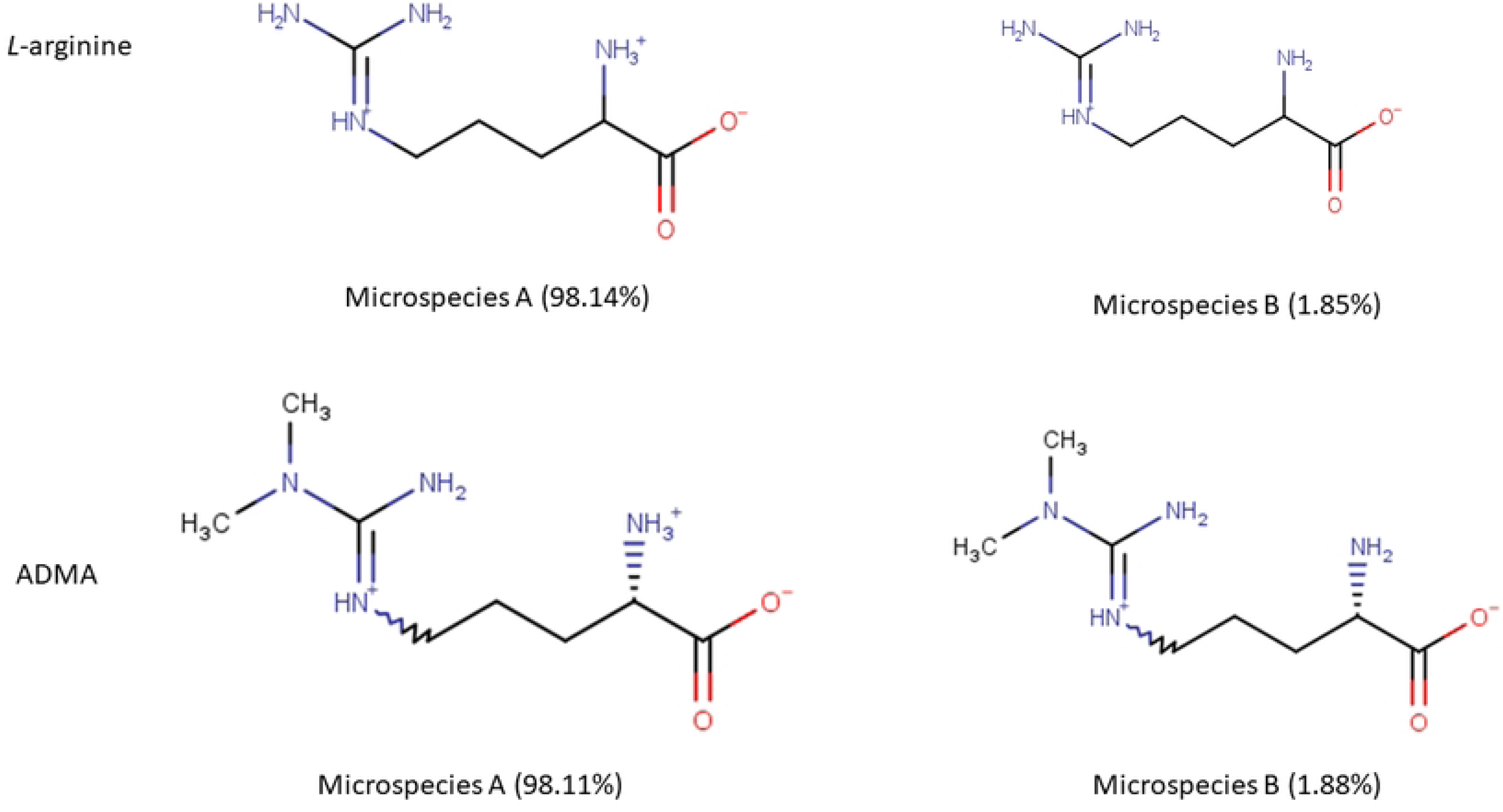
The brain distribution of [^3^H]-arginine and [^14^C]-sucrose as a function of time measured using the *in situ* brain perfusion technique in anaesthetised mice. Uptake is expressed as the percentage ratio of tissue to plasma (mL.100 g^-1^). Each point represents the mean ± SEM of 4-7 animals. K*_in_* and V*_i_* values were determined as the slope and ordinate intercept of the computed regression lines where appropriate and reported in Table 1. Asterisks represent one-tailed, paired Student’s t-tests comparing mean±SEM at each time point, \**p* < 0.05, \*\**p* < 0.01, \*\*\**p* < 0.001 (GraphPad Prism 6.0 for Mac).

**Table 1:** The unidirectional transfer constants (K*_in_*) and initial volume of distribution (V_i_) determined by multiple-time uptake analysis of [^14^C]-sucrose into specific CNS regions. Linear regression analysis produced R^2^ for goodness of fit and *p*-values relating to the deviation of the line slope from zero. There was no significant difference between the [^14^C]-sucrose values reported for the perfusion studies with [^3^H]-arginine (data shown) and for those reported with [^3^H]-ADMA (data not shown).

| Brain Regions | $p$ | $R^2$ | $K_{in}$<br>$\mu\text{l}.\text{min}^{-1}.\text{g}^{-1}$ | $V_i$<br>$\text{mL}.100\text{g}^{-1}$ |
| --- | --- | --- | --- | --- |
| Frontal Cortex | 0.0004 | 0.51 | $0.070 \pm 0.016$ | $1.40 \pm 0.31$ |
| Caudate Nucleus | 0.0005 | 0.50 | $0.067 \pm 0.016$ | $1.19 \pm 1.19$ |
| Occipital Cortex | $<0.0001$ | 0.61 | $0.068 \pm 0.013$ | $1.16 \pm 0.24$ |
| Hippocampus | $<0.0001$ | 0.66 | $0.060 \pm 0.010$ | $0.81 \pm 0.19$ |
| Hypothalamus | 0.0491 | 0.82 | $0.044 \pm 0.023$ | $2.67 \pm 0.43$ |
| Thalamus | 0.0001 | 0.57 | $0.107 \pm 0.022$ | $0.87 \pm 0.41$ |
| Pons | 0.0074 | 0.35 | $0.086 \pm 0.028$ | $2.52 \pm 0.51$ |
| Cerebellum | 0.0026 | 0.40 | $0.058 \pm 0.017$ | $1.66 \pm 0.31$ |
| **Capillary depletion analysis** |  |  |  |  |
| Homogenate | 0.0061 | 0.38 | $0.061 \pm 0.019$ | $1.51 \pm 0.38$ |
| Supernatant | 0.0006 | 0.55 | $0.058 \pm 0.013$ | $0.81 \pm 0.27$ |
| Pellet | 0.0151 | 0.36 | $0.0069 \pm 0.0025$ | $0.21 \pm 0.051$ |

The uptake of [^3^H]-arginine was significantly greater than [^14^C]-sucrose in all brain regions and at all-time points (multiple Student’s paired t-tests, *p* < 0.05 in all cases) (Fig 2). A time-dependent increase in the distribution of [^3^H]-arginine (corrected for [^14^C]-sucrose) was observed in all regions (*e.g.* 28.96±4.03% after 2.5 minutes to 176.78±31.60% after 30 minutes in the frontal cortex) and no regional differences were observed (*p* > 0.05, two-way ANOVA with Tukey’s multiple comparison test for each brain region compared to all others within each time point).

Multiple-time uptake analysis over 30 minutes could not be used to calculate a K*_in_* and V*_i_* for [^3^H]-arginine samples, as equation 3 is only applicable when the amount of substance in V*_i_* is roughly proportional to C*_pl_*. The V*_i_* values determined for [^3^H]-arginine using this method were up to 13-fold higher than that measured for [^14^C]-sucrose space in the same experiment (data not shown). However, single-time uptake analysis (equation 2) to calculate a transfer constant (K*_in_*) for [^3^H]-arginine distribution into all brain regions could be utilized and the values are reported in Table 2. For example, single-time uptake analysis of [^3^H]-arginine at 10 minutes into the frontal cortex, hypothalamus and thalamus revealed a K*_in_* of 5.84±0.86, 7.26±1.76 and 5.87±1.07 μl.min^-1^.g^-1^, respectively (Table 2).

**Table 2:** The *K_in_* values for [^3^H]-arginine and [^3^H]-ADMA values determined by single-time point analysis at a 10-minute perfusion. The R*_Tissue_* values had all been corrected for [^14^C]-sucrose. The rate of uptake of [^3^H]-ADMA is expressed as a percentage of the rate of uptake of [^3^H]-arginine.

| | [$^3\text{H}$]-arginine | | [$^3\text{H}$]-ADMA | | [$^3\text{H}$]-ADMA/<br>[$^3\text{H}$]-arginine |
| --- | --- | --- | --- | --- | --- |
| Brain Region | $K_{in} (\mu\text{l} \cdot \text{min}^{-1} \cdot \text{g}^{-1})$ | n | $K_{in} (\mu\text{l} \cdot \text{min}^{-1} \cdot \text{g}^{-1})$ | n | % |
| Frontal Cortex | $5.84 \pm 0.86$ | 7 | $2.49 \pm 0.35$ | 5 | 42.7 |
| Caudate nucleus | $5.40 \pm 0.67$ | 7 | $2.41 \pm 0.29$ | 5 | 44.6 |
| Occipital Cortex | $4.98 \pm 0.72$ | 7 | $2.25 \pm 0.16$ | 5 | 45.2 |
| Hippocampus | $4.72 \pm 0.56$ | 7 | $2.17 \pm 0.17$ | 5 | 46.1 |
| Hypothalamus | $7.26 \pm 1.76$ | 7 | $3.04 \pm 0.30$ | 5 | 41.9 |
| Thalamus | $5.87 \pm 1.07$ | 7 | $2.10 \pm 0.17$ | 5 | 35.8 |
| Pons | $5.58 \pm 0.97$ | 7 | $2.46 \pm 0.34$ | 5 | 44.2 |
| Cerebellum | $5.10 \pm 0.50$ | 7 | $1.62 \pm 0.19$ | 5 | 31.7 |
| **Capillary depletion analysis** |  |  |  |  |  |
| Homogenate | $3.97 \pm 0.67$ | 7 | $1.27 \pm 0.56$ | 5 | 31.9 |
| Supernatant | $2.35 \pm 0.42$ | 7 | $0.86 \pm 0.16$ | 5 | 36.6 |
| Pellet | $0.70 \pm 0.10$ | 7 | $0.08 \pm 0.20$ | 5 | 11.4 |
| **Circumventricular organs** |  |  |  |  |  |
| CSF | $1.08 \pm 0.24$ | 5 | $2.70 \pm 0.90$ | 5 | 249.5 |
| Choroid plexus | Not applicable | - | $26.09 \pm 4.24$ | 5 | - |
| Pituitary gland | Not applicable | - | $18.28 \pm 3.23$ | 5 | - |
| Pineal gland | Not applicable | - | $19.40 \pm 4.85$ | 5 | - |

Fig 3 confirms the presence of [^3^H]-arginine in the brain homogenate, brain parenchyma-containing supernatant and endothelial cell enriched pellet. There was an increase in the distribution of [^3^H]-arginine in all three capillary depletion samples over time (Fig 3). No differences in the distribution of [^3^H]-arginine between the homogenate and supernatant (containing brain parenchyma) were observed, however both the homogenate and supernatant samples contained a higher distribution than that observed in the endothelial cell-enriched pellet (*p* > 0.01, two-way ANOVA with Tukey’s multiple comparison test for each sample compared to all others).

**Fig 3:**
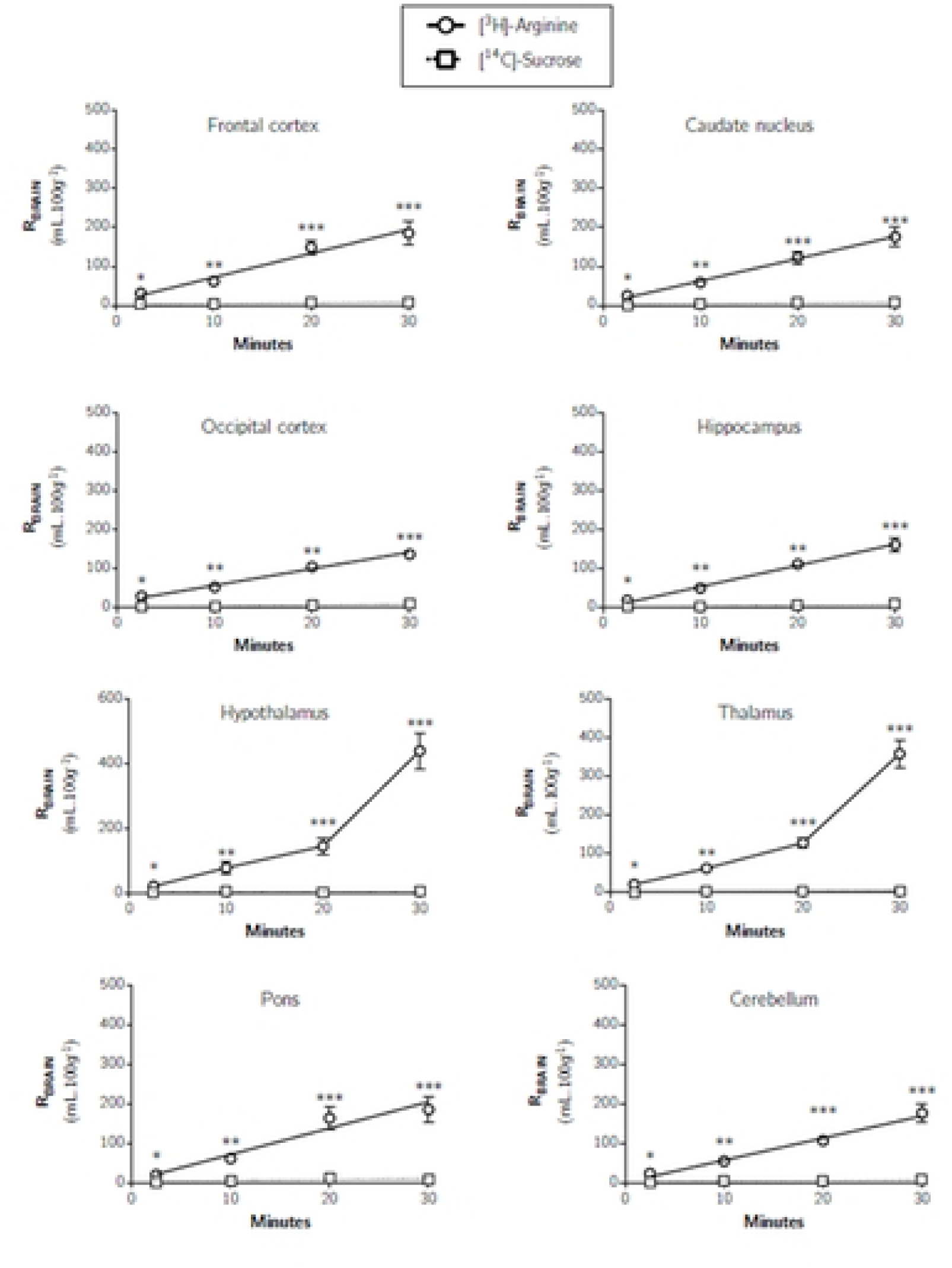
Distribution of [^3^H]-arginine and [^14^C]-sucrose in capillary depletion samples as a function of time. Uptake is expressed as the percentage ratio of tissue to plasma (mL.100 g^-1^). Each point represents the mean ±SEM of 4-7 animals. (GraphPad Prism 6.0 for Mac). Asterisks represent one-tailed, paired Student’s t-tests comparing mean±SEM at each time point, \**p* < 0.05, \*\**p* < 0.01, \*\*\**p* < 0.001.

#### CSF

Fig 4 shows that the [^3^H]-arginine uptake into the CSF was 4.41±1.41% at 2.5 minutes and rose to 120.44±25.08% at 30 minutes (data uncorrected for [^14^C]-sucrose). The K*_in_* determined by single-time uptake analysis for [^3^H]-arginine into the CSF at 10 minutes was 1.08±0.24 μl.min^-1^.g^-1^ (Table 2).

**Fig 4:**
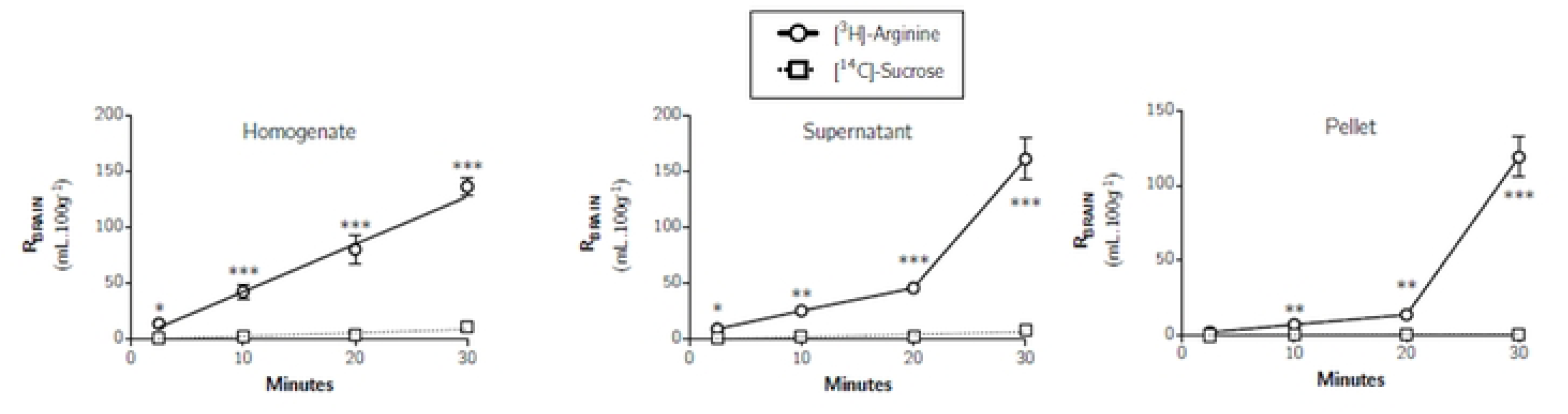
Distribution of [^3^H]-arginine and [^14^C]-sucrose in the CSF, pineal gland, choroid plexus and pituitary gland following *in situ* brain perfusion as a function of time. Uptake is expressed as the percentage ratio of tissue or CSF to plasma (mL.100 g^-1^). Each point represents the mean ± SEM of 4-7 animals. Asterisks represent one-tailed, paired Student’s t-tests comparing mean±SEM at each time point, \**p* < 0.05, \*\**p* < 0.01, \*\*\**p* < 0.001 (GraphPad Prism 6.0 for Mac).

#### Circumventricular Organs

Fig 4 illustrates the distribution of [^14^C]-sucrose and [^3^H]-arginine into the choroid plexus. The [^14^C]-sucrose distribution represents the extracellular space formed between choroid plexus capillary endothelium and epithelium in this highly vascularised tissue. [^3^H]-Arginine accumulation was significantly greater than [^14^C]-sucrose accumulation at all time-points (one-tailed, paired Student’s t-tests comparing mean±SEM at each time point).

Fig 4 illustrates the distribution of [^14^C]-sucrose and [^3^H]-arginine into the pituitary gland and pineal gland. [^3^H]-Arginine accumulation was significantly greater than [^14^C]-sucrose accumulation at all time-points in both tissues (one-tailed, paired Student’s t-tests comparing mean±SEM at each time point). A comparison of the mean [^3^H]-arginine distributions between the pituitary gland and pineal gland revealed that a significant difference is observed between the pituitary gland and pineal gland, but only after a 30-minute perfusion (p<0.001, two way ANOVA with Tukey’s multiple comparison test for each sample compared to all others within each time point). Single-time uptake analysis could not be used to determine a K*_in_* for [^3^H]-arginine transfer into the choroid plexus, pituitary gland and pineal gland as the [^3^H]-arginine concentration in these tissues was higher than the concentration in the plasma (Table 2). Single-time uptake analysis to calculate a transfer constant can only be applied if entry of the test solute into the CNS is proportional to its plasma concentration and the concentration in the CNS is less than the concentration in the plasma and efflux (CNS to blood) is much smaller than influx (blood to CNS) of the test solute and therefore can be ignored (22).

### ADMA

#### Brain Distribution

Fig 5 shows the distribution (R_Brain_) of [^14^C]-sucrose and [^3^H]-ADMA into the frontal cortex, caudate nucleus, occipital cortex, hippocampus, hypothalamus, thalamus, pons and cerebellum over time. The uptake of [^14^C]-sucrose was low in all brain regions confirming the integrity of the BBB. The uptake of [^14^C]-sucrose was significantly lower than [^3^H]-ADMA in all brain regions and at all-time points (multiple Student’s paired t-tests, *p* < 0.05 in all cases).

**Fig 5:**
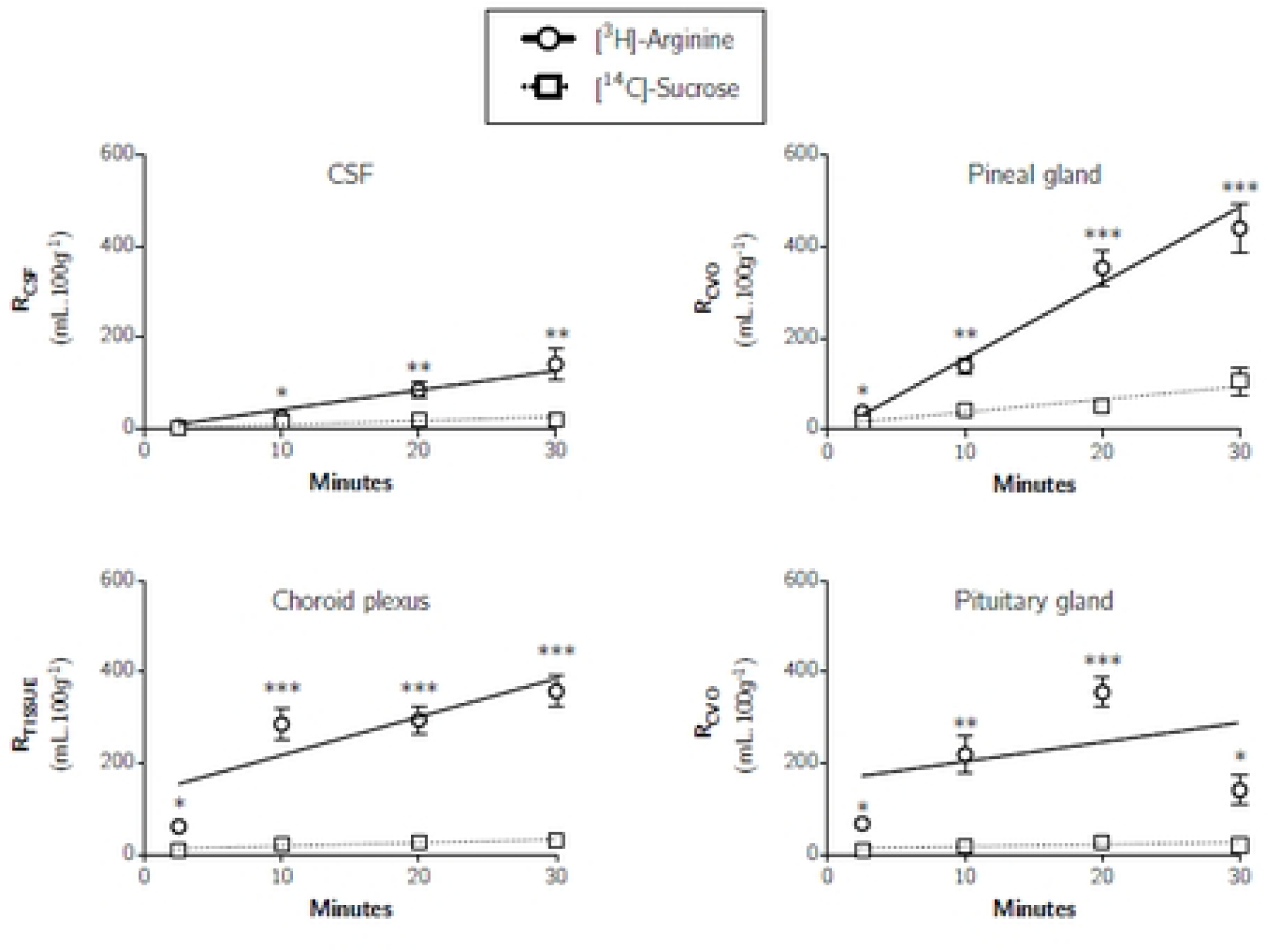
Brain distribution of [^3^H]-ADMA and [^14^C]-sucrose as a function of time. Uptake is expressed as the percentage ratio of tissue to plasma (mL.100 g^-1^). Each point represents the mean ± SEM of 5 animals. One-tailed, paired Student’s t-tests comparing mean±SEM at each time point, \**p* < 0.05, \*\**p* < 0.01 (GraphPad Prism 6.0 for Mac).

A time-dependent increase in the distribution of [^3^H]-ADMA (corrected for [^14^C]sucrose) was observed in all regions up to 20 minutes (*e.g.* 9.86±1.43% after 2.5 minutes to 27.64±4.30% after 20 minutes in the frontal cortex), however this was followed by a decrease in the distribution of [^3^H]-ADMA at 30 minutes in all regions (*e.g.* 10.41±2.77% in the frontal cortex). No regional differences were observed (p > 0.05, two-way ANOVA with Tukey’s multiple comparison test for each brain region compared to all others within each time point). Due to a departure of linearity of the experimental points, multiple time uptake analysis (i.e. equation 3) could not be used to calculate a K*_in_* for [^3^H]-ADMA, however, single time uptake analysis at 10 minutes was acceptable and revealed a K*_in_* of 2.49±0.35 μl.min^-1^.g^-1^ for [^3^H]-ADMA distribution into the frontal cortex (Table 2). There was no significant difference of the K*_in_* obtained from the eight brain regions sampled (one-way ANOVA with Dunnett’s multiple comparisons of means, *p* > 0.05).

Fig 6 confirms the presence of [^3^H]-ADMA in the brain parenchyma-containing supernatant and endothelial cell enriched pellet. This suggests that [^3^H]-ADMA can cross the BBB and enter the brain tissue. A very similar pattern of a time-dependent peak in the distribution of [^3^H]-ADMA after perfusing for 20 minutes was observed in whole brain homogenate and brain parenchyma (supernatant) following capillary depletion (Fig 6) to that observed in the brain regions (Fig 5). Interestingly, the peak in [^3^H]-ADMA distribution in the endothelial cell-enriched pellet following capillary depletion was observed earlier, after only 10 minutes of perfusion.

**Fig 6:**
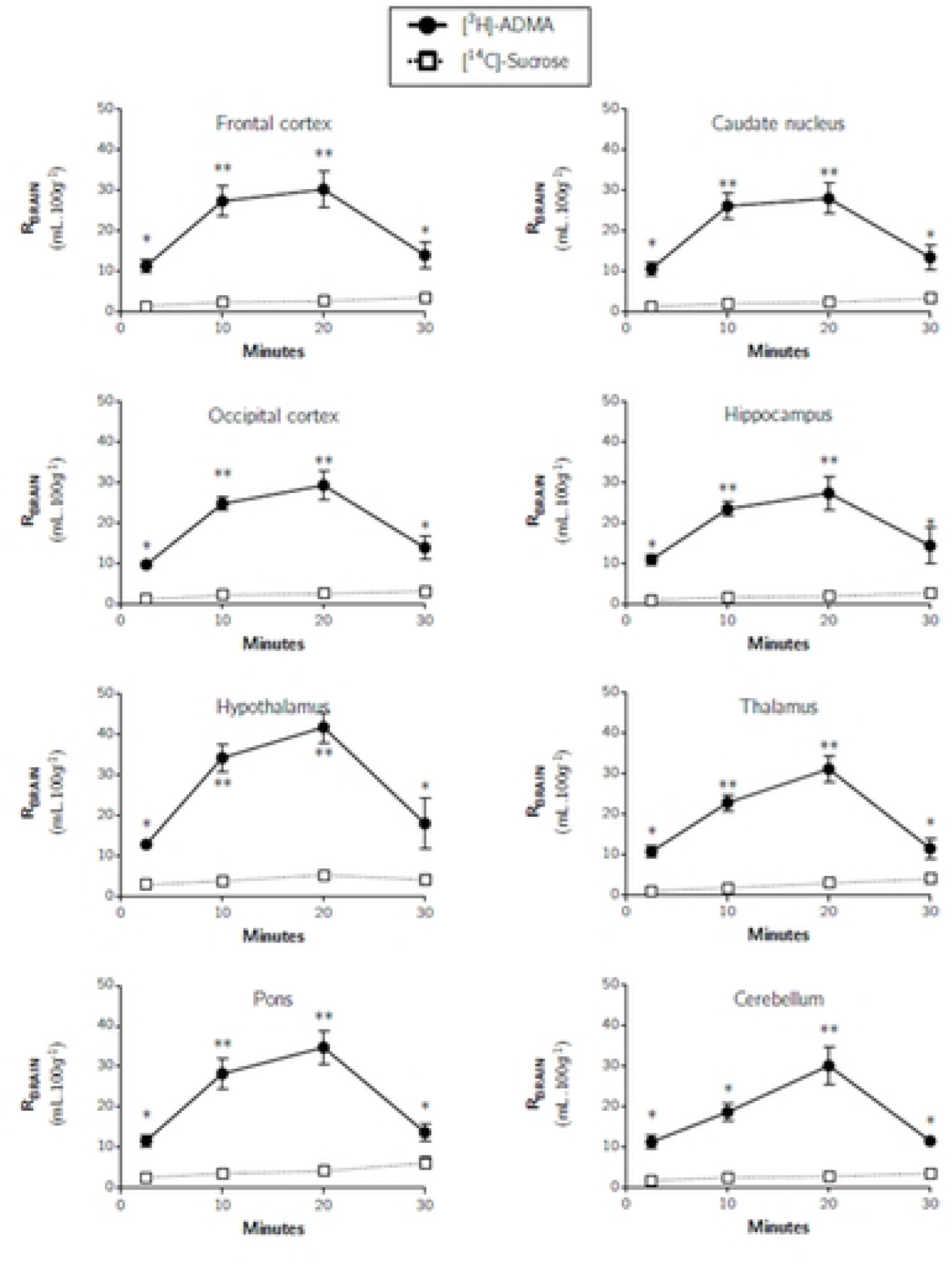
Distribution of [^3^H]-ADMA and [^14^C]-sucrose in capillary depletion samples as a function of time. Uptake is expressed as the percentage ratio of tissue to plasma (mL.100 g^-1^). Each point represents the mean ± SEM of 5 animals. One-tailed, paired Student’s t-tests comparing mean±SEM at each time point, \**p* < 0.05, \*\**p* < 0.01 (GraphPad Prism 6.0 for Mac).

### CSF

Fig 7 shows that [^3^H]-ADMA could be detected in the CSF at 2.5 minutes being approximately 4%. Interestingly, the CSF distribution of [^3^H]-ADMA was like [^14^C]-sucrose at 2.5 and 30 minutes, but significantly higher at 10 and 20 minutes (*p*<0.05: One-tailed, paired Student’s t-test comparing mean±SEM at each time point). The K*_in_* value determined by single-time uptake analysis at 10 minutes for [^3^H]-ADMA in the CSF was not significantly different to that in any of the brain regions samples (Table 2; one-way ANOVA with Dunnett’s multiple comparisons of means, *p*>0.05).

**Fig 7:**
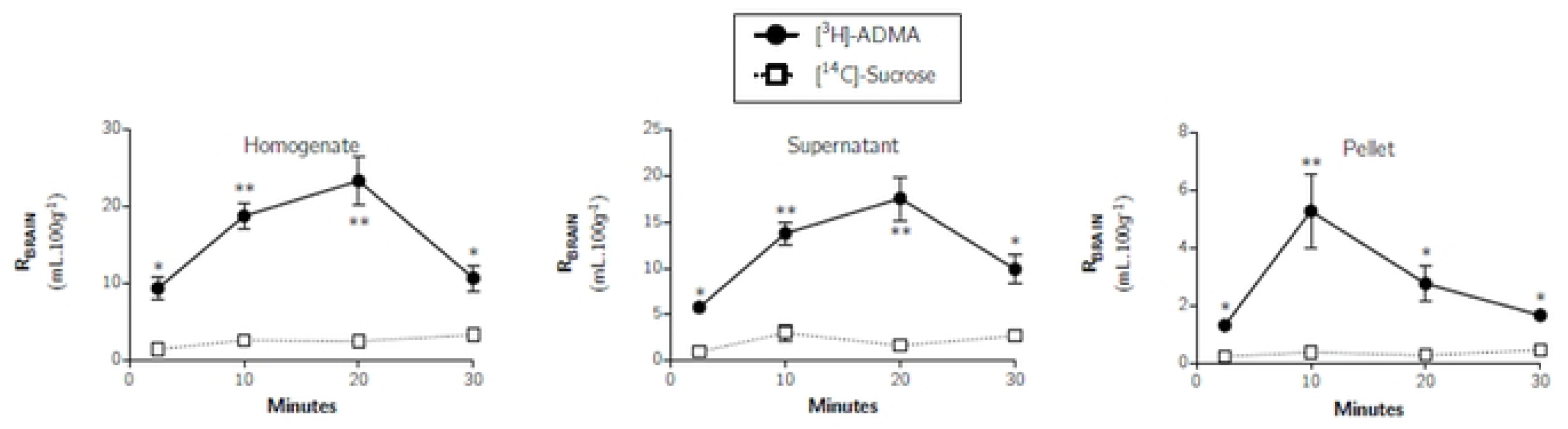
Distribution of [^3^H]-ADMA and [^14^C]-sucrose in the CSF, pineal gland, choroid plexus and pituitary gland following *in situ* brain perfusion as a function of time. Uptake is expressed as the percentage ratio of tissue or CSF to plasma (mL.100 g^-1^). Each point represents the mean±SEM of 5 animals. One-tailed, paired Student’s t-tests comparing mean±SEM at each time point, \**p*< 0.05, ** *p*< 0.01, *** *p*< 0.001 (GraphPad Prism 6.0 for Mac).

#### Circumventricular organs

Fig 7 illustrates the distribution of [^14^C]-sucrose and [^3^H]-ADMA into the choroid plexus. The [^14^C]-sucrose distribution represents the extracellular space in this highly vascularised tissue. [^3^H]-ADMA accumulation was significantly greater than [^14^C]-sucrose accumulation at 2.5, 10 and 20 minutes. Importantly, the distribution of [^3^H]-ADMA into the choroid plexus reached a peak at 10 minutes of approximately 240% and then started to decrease until it reached approximately 40% at 30 minutes. A similar time-dependent increase and then decrease in the distribution of [^3^H]-ADMA was also observed in the pituitary gland and pineal gland (Fig 7). The mean [^3^H]-ADMA distribution in the choroid plexus, pituitary gland and pineal gland was not significantly different at any time point (uncorrected values, two-way ANOVA with Tukey’s multiple comparison test for each sample compared to all others within each time point). In addition, the *K_in_* values for the choroid plexus, pituitary gland and pineal gland, were not significantly different from each other, but were significantly higher than all other samples (*i.e.* brain regions and CSF) (one-way ANOVA with Dunnett’s multiple comparisons of means, *p*< 0.05).

### Comparison of [^3^H]-arginine and [^3^H]-ADMA

The uptake of [^3^H]-arginine, [^3^H]-ADMA and [^14^C]-sucrose as a function of time is compared in S1-S3 Fig. The uptake of [^3^H]-arginine into the eight brain regions was significantly higher than that of [^3^H]-ADMA (*p* < 0.05) at all time points (Student’s unpaired, two-tailed t-tests were used for the comparison of the two means) (S1 Fig). The rate of uptake of [^3^H]-ADMA ranged from 31.7-46.1% of the rate of uptake of [^3^H]-arginine into the different brain regions at 10 minutes (Table 2).

In the whole brain homogenate, and resulting supernatant following capillary depletion, [^3^H]-arginine distribution was significantly higher than [^3^H]-ADMA at 10-, 20- and 30-minute time points (*p* < 0.05) (S2 Fig). In the endothelial cell-enriched pellet resulting from capillary depletion, [^3^H]-arginine distribution was only significantly higher than [^3^H]-ADMA at the 20- and 30-minute time points (*p* < 0.01). The rate of uptake of [^3^H]-ADMA ranged from 11.4-36.6 % of the rate of uptake of [^3^H]-arginine into the different samples at 10 minutes (Table 2).

In the case of the CSF, pituitary gland, pineal gland and choroid plexus, [^3^H]-arginine distribution was significantly higher than [^3^H]-ADMA at the 20- and 30-minute time points (p < 0.05) (S3 Fig). The rate of uptake of [^3^H]-ADMA ranged from 90.4-204.0 % of the rate of uptake of [^3^H]-arginine into the different samples at 10 minutes (Table 2).

### Self-inhibition experiments

Fig 8 shows the effect of 100μM unlabelled arginine on the uptake of [^3^H]-arginine into all brain regions after a 10-minute perfusion. These data indicate that [^3^H]-arginine uptake into all eight brain regions is markedly self-inhibited by an average of approximately 67% (*p* < 0.05, unpaired, one-tailed Student’s t-test comparing means). S4 Fig shows the effect of 100μM unlabelled arginine on the distribution of [^3^H]-arginine into capillary depletion samples. These data indicate that [^3^H]-arginine uptake is inhibited by an average of approximately 56% (*p* < 0.05, unpaired, one-tailed Student’s t-test comparing means). S5 Fig also shows the effect of 100μM on the unlabelled *L*-arginine on the distribution of [^3^H]-arginine into the CSF. These data indicate that [^3^H]-arginine uptake into the CSF is inhibited by 62.2% (*p* < 0.01, unpaired, one-tailed Student’s t-test comparing means). S5 Fig also shows the inhibitory effect of an excess of unlabelled arginine on the distribution of [^3^H]-arginine into choroid plexus (57.3%; *p*<0.001), pineal gland (39.4%; *p*<0.05) and pituitary gland (48.1%; *p*<0.05) (unpaired, one-tailed Student’s t-test comparing the means).

**Fig 8:**
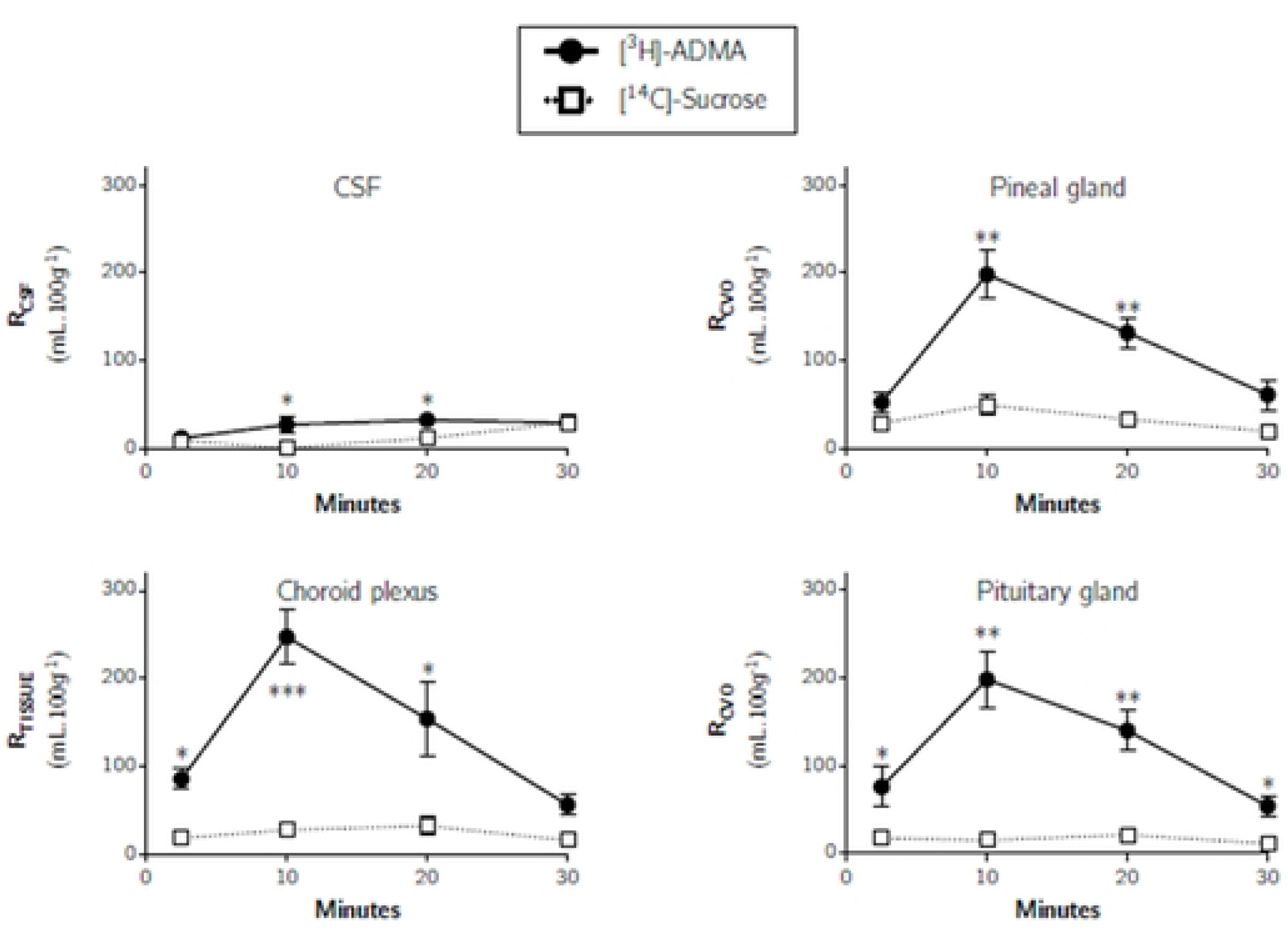
The effect of 100μM unlabelled *L*-arginine on the uptake of [^3^H]-arginine in the brain. Uptake is expressed as the percentage ratio of tissue to plasma (mL.100 g^-1^) and is corrected for [^14^C]-sucrose (vascular space). Perfusion time is 10 minutes. Each bar represents the mean ± SEM of 6-7 animals (GraphPad Prism 6.0 for Mac). \**p* < 0.05, \*\**p* < 0.01, \*\*\**p* < 0.001.

The uptake of [^3^H]-ADMA is also significantly self-inhibited by 100 μM unlabelled ADMA in all brain regions after a 10-minute perfusion (Fig 9). The uptake of [^3^H]-ADMA being significantly decreased by 60.3 to 74.3 % when unlabelled ADMA was present. The same phenomenon was observed in capillary depletion samples (reduction 77.1 to 95.2%), CSF (reduction 89.1%) and circumventricular organs (reduction 68.0 - 82.4%) (S6 Fig and S7 Fig).

**Fig 9:**
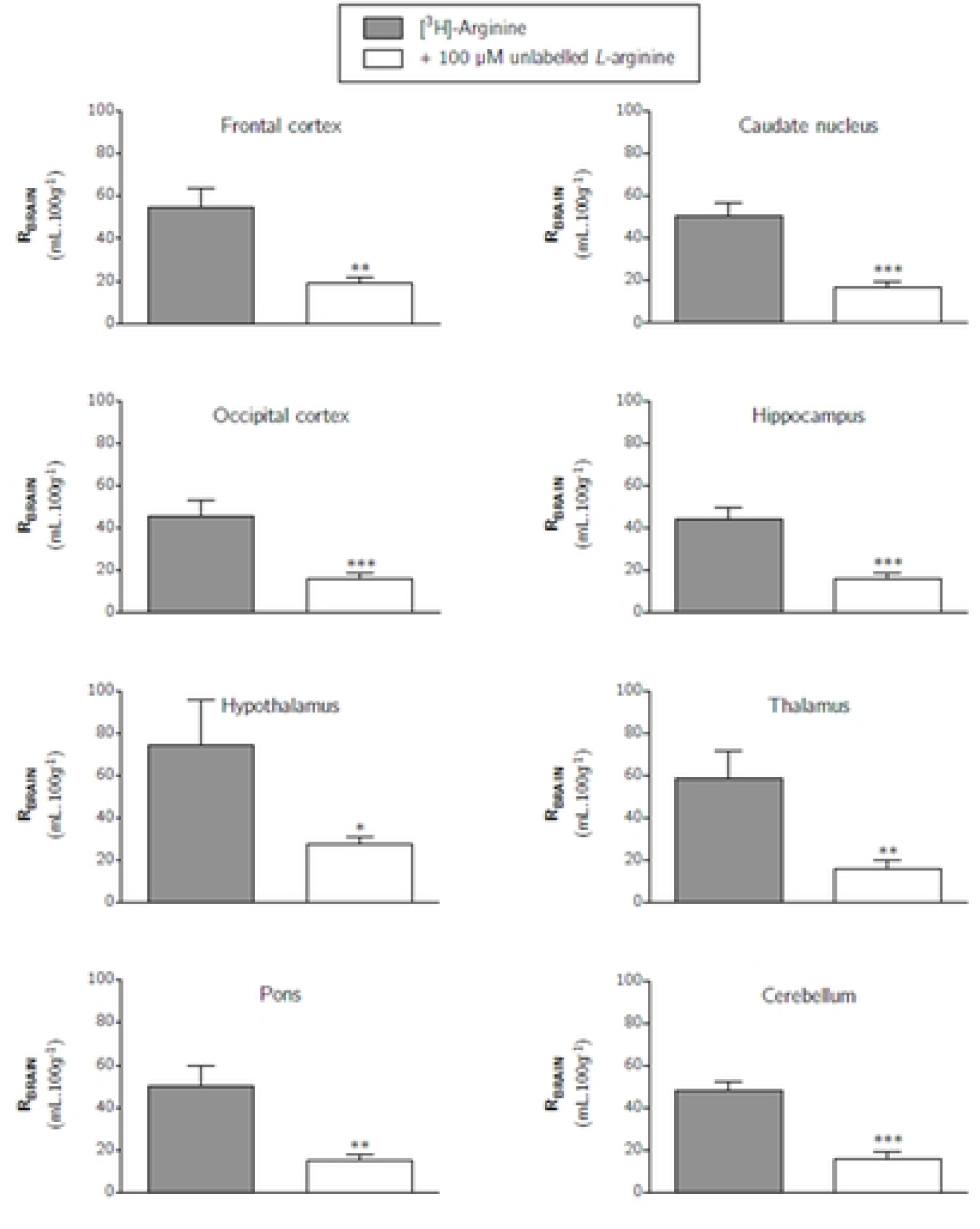
The effect of 100μM unlabelled ADMA on the uptake of [^3^H]-ADMA in the brain. Uptake is expressed as the percentage ratio of tissue to plasma (mL.100 g^-1^) and is corrected for [^14^C]-sucrose (vascular space). Perfusion time is 10 minutes. Each bar represents the mean ± SEM of 4-5 animals (GraphPad Prism 6.0 for Mac). \*\*\**p* < 0.001.

### Cross-inhibition experiments

To determine the effect of ADMA on the transport of [^3^H]-*L*-arginine across the blood-brain and blood-CSF barriers, [^3^H]-arginine was co-perfused with different concentrations of unlabelled ADMA. Concentrations of ADMA were specifically selected to mimic plasma concentrations under normal conditions (0.5 μM), pathophysiological conditions (3.0 μM), and then supraphysiological concentrations (10, 100 and 500 μM). Fig 10 shows the effect of an excess of unlabelled ADMA on the uptake of [^3^H]-arginine into all brain regions. These data indicate that [^3^H]-arginine uptake is only significantly inhibited by the highest concentration of unlabelled ADMA, which was 500 μM. [^3^H]-arginine uptake being significantly inhibited by approximately 70% into each of the brain regions (one-way ANOVA with Dunnett’s post-hoc test comparing means to control, *p* <0.05). Capillary depletion analysis of remaining whole brain tissue revealed that distribution of [^3^H]-arginine in only the whole brain homogenate and resulting supernatant (brain parenchyma) was inhibited by 500 μM unlabelled ADMA by 65.1% and 72.0%, respectively (Fig 11, *p* < 0.05). However, the distribution of [^3^H]-arginine into the cerebral capillary endothelial cells (i.e. pellet) was not affected by unlabelled ADMA even at the highest concentration of 500 μM (Fig 11). The distribution of [^3^H]-arginine in the CSF, pineal gland and pituitary gland was also not affected by any of the concentrations of unlabelled ADMA included in artificial plasma (Fig 12). However, 500 μM unlabelled ADMA inhibited the distribution of [^3^H]-arginine in the choroid plexus by 72.6% (*p* < 0.01).

**Fig 10:**
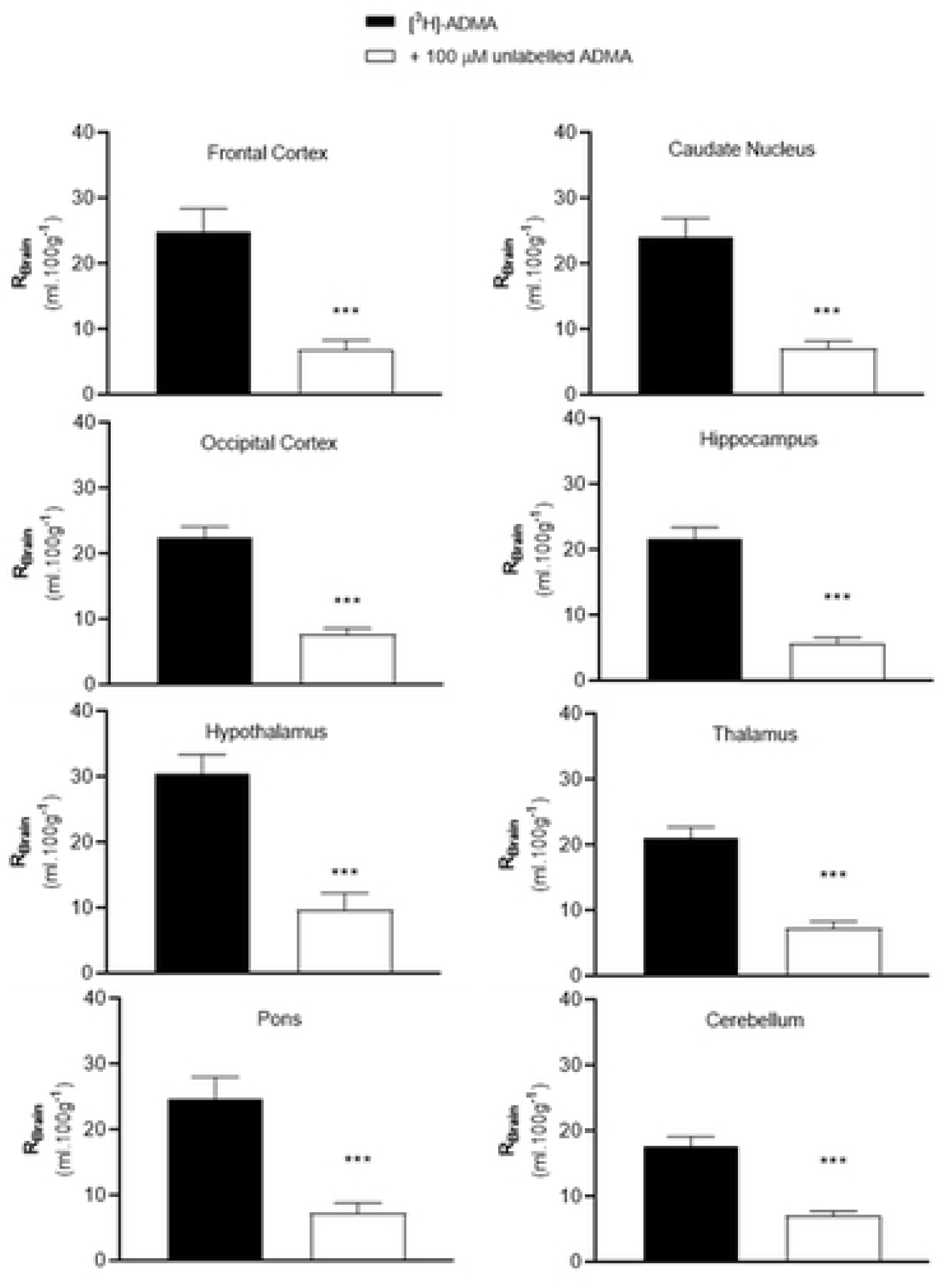
The effect of unlabelled ADMA on the regional brain uptake of [^3^H]-arginine (10 minute perfusion). Uptake is expressed as the percentage ratio of tissue to plasma (mL.100 g^-1^) and is corrected for [^14^C]-sucrose (vascular space). Perfusion time is 10 minutes. Each bar represents the mean ± SEM of 4-7 animals. Asterisks represent one-way ANOVA with Dunnett’s post-hoc tests comparing mean±SEM to control within each region/sample, \**p* <0.05, \*\**p* < 0.01 (GraphPad Prism 6.0 for Mac).

**Fig 11:**
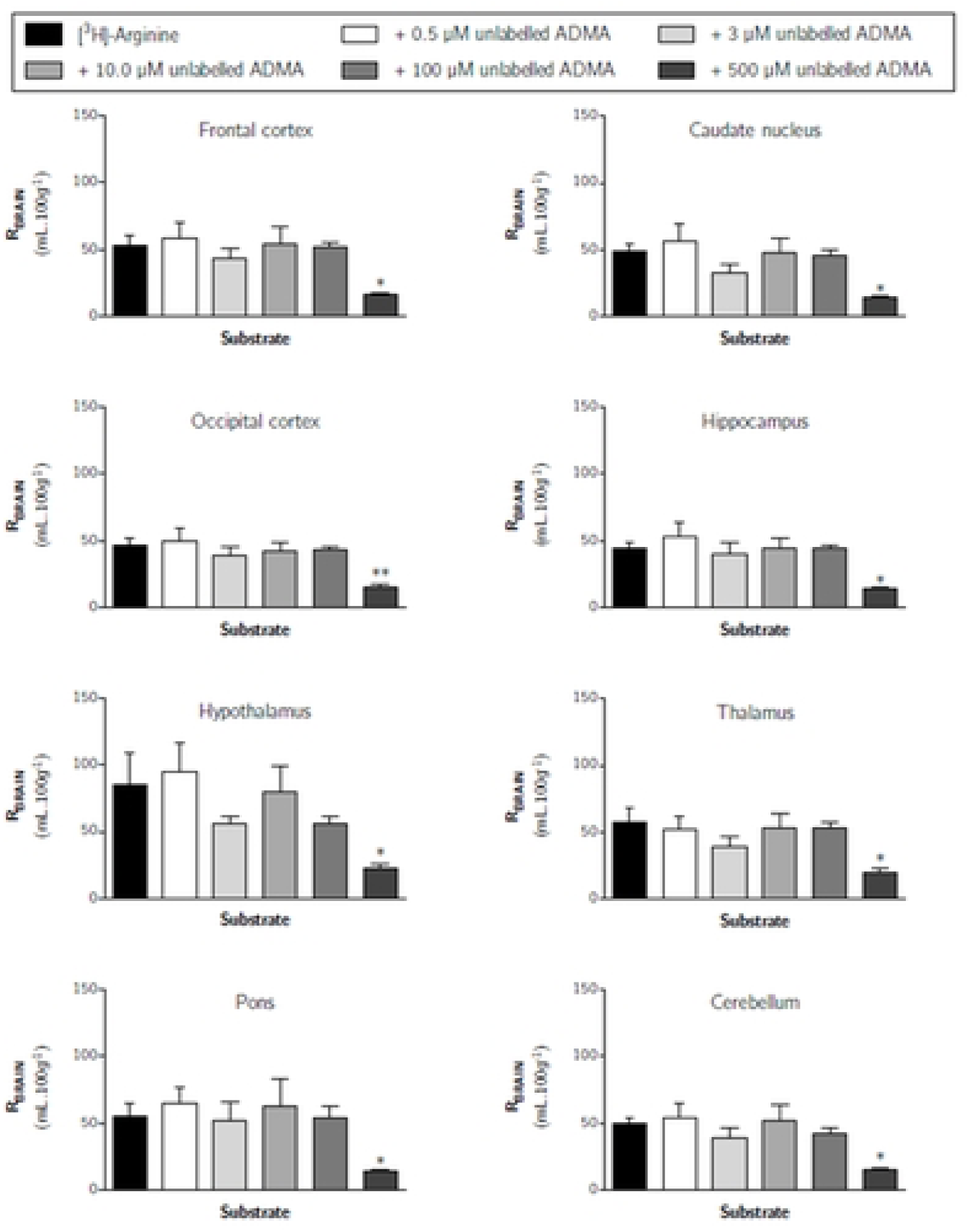
The effect of unlabelled ADMA on the distribution of [^3^H]-arginine in capillary depletion samples (10 minute perfusion). Uptake is expressed as the percentage ratio of tissue to plasma (mL.100 g^-1^) and is corrected for [^14^C]-sucrose (vascular space). Perfusion time is 10 minutes. Each bar represents the mean ± SEM of 4-7 animals. Asterisks represent one-way ANOVA with Dunnett’s post-hoc tests comparing mean±SEM to control within each region/sample, \**p* < 0.05 (GraphPad Prism 6.0 for Mac).

**Fig 12:**
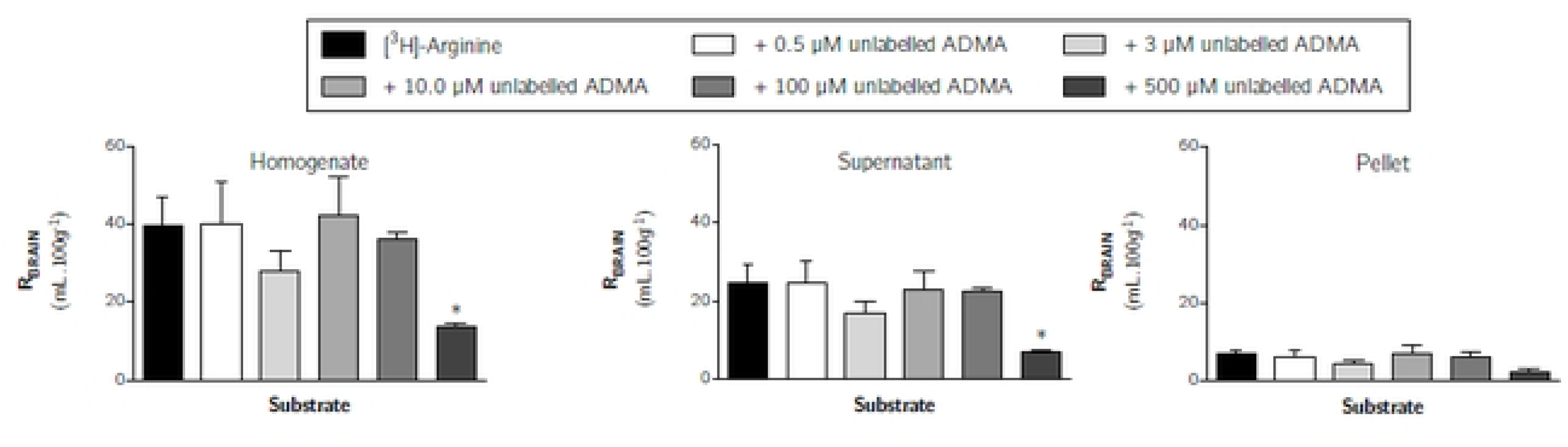
The effect of unlabelled ADMA on the distribution of [^3^H]-arginine in CSF, choroid plexus and CVOs (10 minute perfusion). Uptake is expressed as the percentage ratio of tissue or CSF to plasma (mL.100 g^-1^). Perfusion time is 10 minutes. Each bar represents the mean ± SEM of 4-7 animals. Asterisks represent one-way ANOVA with Dunnett’s post-hoc tests comparing mean±SEM to control within each region/sample, \*\**p* < 0.01 (GraphPad Prism 6.0 for Mac).

Whilst 100 μM unlabelled ADMA did not inhibit the distribution of [^3^H]-arginine in any of the samples analysed, 100 μM *L*-arginine was sufficient to significantly inhibit distribution of [^3^H]-ADMA (S8-S10 Figs). S8 Fig shows the effect of an excess of unlabelled ADMA on the uptake of [^3^H]-arginine into all brain regions. These data indicate that [^3^H]-ADMA uptake is inhibited by 100 μM unlabelled *L*-arginine by up to 80.4% (one-tailed unpaired Student’s t-test comparing means, p < 0.001). This trend was also mirrored in samples from capillary depletion analysis where 100 μM *L*-arginine inhibited [^3^H]-ADMA distribution in all samples by up to 80.0% (S9 Fig, *p* < 0.01).

S10 Fig shows the effect of an excess of unlabelled *L*-arginine on the distribution of [^3^H]-ADMA into the CSF, pineal gland, choroid plexus and pituitary gland. These data indicate that [^3^H]-ADMA uptake is markedly inhibited by up to 78.6% (*p* < 0.05, unpaired, one-tailed Student’s t-test comparing means). The inclusion of unlabelled *L*- arginine however had no effect on the distribution of [^3^H]-ADMA in the CSF.

The degree of inhibition of [^3^H]-ADMA distribution exerted by the inclusion of either 100 μM unlabelled *L*-arginine (cross-competition) and 100 μM unlabelled ADMA (self-inhibition) appears to be similar (S11 Fig). This contrasts with the difference in sensitivity observed when [^3^H]-arginine distribution into the CNS was examined in the presence of 100 μM unlabelled arginine (significantly reduced) or 100 μM unlabelled ADMA (no change) (S11 Fig).

## DISCUSSION

Circulating levels of the cationic amino acid, *L*-arginine, and its endogenously produced homologue, ADMA, are in a delicately poised equilibrium. As arginine is a NOS substrate and ADMA is a NOS inhibitor, dysregulation of this balance has well-documented implications in cardiovascular disease and brain conditions, as it affects NO production. In fact, impaired NO bioavailability represents the central feature of endothelial dysfunction, which is found in many diseases for example Alzheimer’s disease (3). The principal aim of our study is to explore ADMA and *L*-arginine transport at the blood-brain and blood-CSF barriers and discuss how it could be manipulated by arginine supplementation to treat NO dysregulation.

The first part of our study utilized a specialist chemical property database to explore the physicochemical characteristics of *L*-arginine and ADMA (15). This stated that *L*-arginine and ADMA had a molecular weight of 174.2 and 202.26 g/mol respectively and had a gross charge at physiological pH of +0.981. Interestingly, both *L*-arginine and ADMA exist as two microspecies at physiological pH. The major microspecies of both amino acids is a cationic (tripolar) amino acid (∼98.1%) and the other microspecies is a zwitterion (dipolar) amino acid (∼1.8%) (Fig 1). *L*-arginine has a lower lipophilicity than ADMA (predicted log D at pH 7.4 of −4.77 versus −3.99). This relationship was confirmed using octanol:saline partition coefficient measures.

The second part of our study compared the ability of [^3^H]-arginine and [^3^H]-ADMA to cross the mouse blood-CNS barriers and accumulate within the CNS. The integrity of the blood-CNS barriers was confirmed in all experiments by means of the marker molecule, [^14^C]-sucrose.

Multiple-time uptake studies detected [^3^H]-arginine in all brain regions (including frontal cortex, caudate nucleus, occipital cortex, hippocampus, hypothalamus, thalamus, pons and cerebellum) and CVOs (including the choroid plexus, pituitary gland and pineal gland) at significantly higher concentrations than [^14^C]-sucrose. The uptake of [^3^H]-arginine into the capillary endothelial cell enriched pellet and the CSF was also significantly higher than [^14^C]-sucrose at all time points except 2.5 minutes where there was no difference. These results would indicate that [^3^H]-arginine can cross the cerebral capillary endothelium (i.e. the site of the BBB) to reach brain cells and that [^3^H]-arginine can cross the blood-CSF barrier (at the choroid plexuses) to reach the CSF.

While *in situ* brain/choroid plexus perfusion has previously been used to study [^3^H]-arginine uptake into the rat brain (27), no absolute values were published, and thus there are no published values to compare to the results described in our study. In our study, after 30 minutes, brain uptake of [^3^H]-arginine as a percentage of perfused plasma levels reached approximately 185.1% in the pons, and was similarly high in the majority of other brain regions, including the frontal cortex, caudate nucleus, occipital cortex, hippocampus and cerebellum (Fig 2). Thus after 30 minutes it appears that [^3^H]-arginine is sequestered within the brain parenchyma at levels greater than the artificial plasma levels (R_Brain_ > 100%; [^3^H]-arginine artificial plasma concentration of 11.6 nM). One may predict that this would create a concentration gradient favouring flux of [^3^H]-arginine from the brain back into the plasma, at least under the experimental conditions presented here. However, this is unlikely to occur *in vivo* under normal conditions as the concentration of *L*-arginine in plasma is typically 100 μM in humans (28) and 140 μM in mice (29), while brain concentrations in both mice and humans are typically much lower at approximately 0.2-0.3 μM (30) (31). This would create a concentration gradient that overwhelmingly favours facilitated transport of *L*-arginine from plasma into the brain.

Multiple-time studies revealed that [^3^H]-ADMA uptake into all brain regions, capillary depletion samples (homogenate, supernatant and pellet) and pituitary gland were significantly higher than [^14^C]-sucrose at all time points (Fig 5, Fig 6 and Fig 7). The uptake of [^3^H]-ADMA into the choroid plexus was also significantly higher than [^14^C]-sucrose at all time points except 30 minutes where there was no difference. The uptake of [^3^H]-ADMA into the pineal gland and the CSF was also significantly higher than [^14^C]-sucrose at all time points except 2.5 and 30 minutes where there was also no difference. These results would indicate that [^3^H]-ADMA can cross the cerebral capillary endothelium (i.e. the site of the BBB) to reach brain cells and that [^3^H]-ADMA can cross the blood-CSF barrier at the choroid plexuses to reach the CSF.

Another important point is that transport of [^3^H]-ADMA appears to be bi-phasic in nature, with a peak in accumulation being reached at 10 to 20 minutes before decreasing at 30 minutes in all samples. It is possible that these results are related to the effect of ADMA on NO production, which would cause the cerebral blood vessels to vasoconstrict, reducing blood flow. This would decrease [^3^H]-ADMA delivery to the CNS. However, the [^14^C]-sucrose cerebrovascular space measured in the [^3^H]-ADMA study (Fig 5, Fig 6 and Fig 7) was not statistically different to that in the [^3^H]-arginine study (Fig 2, Fig 3 and Fig 4) so the inhibition of eNOS by ADMA is unlikely to generate the results observed. As the transport of the radiolabelled solute of interest back from the brain to the blood can be detected by a loss of linearity of the experimental points (21), the results of this present study indicate a significant CNS-to-blood efflux of [^3^H]-ADMA.

Interestingly, it has been shown that ADMA produced by one cell (e.g. an endothelial cell) can inhibit nitric oxide synthesis in a neighbouring cell (e.g. a macrophage)(32).

Transporters which can remove ADMA from cells include system y^+^ and system y^+^,L (12) (13). Both these transport systems are expressed at the BBB (27,33–35) and the system y^+^ transporter protein, CAT1, is also expressed at the choroid plexus (27)(36). These facts provide further evidence that ADMA could be transferred from the CNS to the blood across the BBB and blood-CSF barrier. It is possible that the activity of these removal mechanism(s) become apparent above a threshold ADMA concentration possibly due to transporter saturation hence the biphasic nature of the data shown in Fig 5, Fig 6 and Fig 7. This would suggest that multiple saturable transport mechanisms with different kinetics are involved in ADMA transfer across cell membranes. In fact, the results so far described would indicate that [^3^H]-ADMA can cross the cerebral capillary endothelium (i.e. the site of the BBB) and the blood-CSF barrier in both directions (i.e. blood-to-CNS and CNS-to-blood).

During the experimental period when unidirectional transfer constants (K*_in_*) can be calculated from multiple time uptake data using equation 3, the amount of test substance in V*_j_* is roughly proportional to C_pl_ and the test substance moves unidirectionally from plasma into brain tissue (26). Consequently, multiple-time uptake analysis could not be used to calculate kinetic constants for either [^3^H]-arginine, as the former condition was not met, nor for [^3^H]-ADMA, as the latter condition was not met. Therefore, to calculate a transfer constant, inhibitor studies at a perfusion time of 10 minutes were performed (Fig 8, Fig 9 and Fig 10) followed by single time uptake analysis (Table 2). Single-time uptake analysis to calculate a transfer constant using equation 2 can be applied if entry of the test solute into the CNS is proportional to its plasma concentration, the concentration in the CNS is less than the concentration in the plasma and efflux (CNS-to-blood) is much smaller than influx (blood-to-CNS) of the test solute and therefore can be ignored (22). Therefore, a transfer constant could be calculated for [^3^H]-arginine and [^3^H]-ADMA distribution into most CNS regions at a perfusion time of 10 minutes (Table 2). Although, it was not possible to determine a transfer constant by single-time uptake analysis for [^3^H]-arginine distribution into the choroid plexus, pineal gland and pituitary gland, as its concentration in the CNS was more than the plasma (Fig 4).

Of the two radiolabelled cationic amino acids studied, [^3^H]-arginine demonstrated the greatest ability (more than two-fold in most cases) to cross the BBB and accumulate within all eight brain regions and the capillary depletion samples (S1 Fig and S2 Fig; Table 2). Both *L*-arginine and ADMA have the same gross charge of +0.981 and a major microspecies with a single positive charge at physiological pH (Fig 1). As arginine has a higher hydrophilicity when compared to ADMA and is of similar hydrophilicity to sucrose, the greater ability of arginine to accumulate within all regions of the brain may be related to the use of specific transporters. These transporters may aid the movement of [^3^H]-arginine into the brain and / or aid the removal of [^3^H]-ADMA out of the CNS.

Our self-inhibition studies using the *in situ* brain/choroid plexus perfusion technique and 100 μM unlabelled arginine at 10 minutes caused a reduction in the uptake of [^3^H]-arginine (11.6 nM) by approximately 39.4-73.0 % into all the CNS samples measured. This would indicate the use of transporters to transfer [^3^H]-arginine across the mouse blood-brain and blood-CSF barriers. The results of our study would also confirm that there are (at the very least) transporters for [^3^H]-arginine on the luminal membrane of the cerebral capillary endothelium (pellet sample) and the blood-side of the choroid plexus.

Saturable transport of radiolabelled *L*-arginine has previously been observed across the rat BBB using the *in situ* brain perfusion technique (37) (27) and at the blood-CSF barrier using the isolated perfused sheep choroid plexus technique (38) (39). Although we did not determine the half-saturation constant (K_m_) of [^3^H]-arginine transport across the blood-CNS interfaces in our mouse study, our data would align with the half-saturation constant (K_m_) for arginine at the BBB previously determined using a similar method in rats (56 ± 9 μM) and the brain uptake index method in rats (40 ± 24 μM (±standard deviation))(37) and the blood-side (basolateral membrane) of the choroid plexus epithelium determined using the isolated perfused choroid plexus of the sheep (25.4 ± 5.1 μM) (27) (40).

Kinetic studies in different types of human and animal cells have indicated that sodium-independent influx of arginine at physiological concentrations occurs predominately by one saturable transport system and has extracellular K_m_ values ranging from 25 to 200 μM (41)(42)(43) (44).

Interestingly, the cationic amino acids transport system, system-y^+^, has been detected at both luminal and abluminal membranes in brain capillary endothelial cells, with prevalence at the abluminal membrane (35). In addition, the half-saturation constants (K_m_) for the y^+^-system expressed in Xenopus oocytes and human fibroblasts is 77 ± 2 μM and 40 ± 0.05 μM, respectively (27)(45)(43). It is therefore plausible that the transporters for arginine detected at the blood-CNS interfaces in our study are system-y^+^ transporters. We have previously identified the presence of the y^+^-system protein, CAT1, in BBB cells (hCMEC/D3) using Western blotting and immunofluorescence (14)(46) and CAT 1 is expressed at the blood-CSF barrier specifically the choroid plexus (27)(36).

Saturable transport of [^3^H]-ADMA into cultured human cerebral capillary endothelial cells has previously been demonstrated by our group and was suggested to be due to system y^+^ - activity (14). In our present *in situ* mouse brain perfusion study, self-inhibition studies using 100 μM unlabelled ADMA at 10 minutes reduced the uptake of [^3^H]-ADMA (62.5 nM) by approximately 60.3-95.2 % into all the CNS samples measured. This indicates the use of transporters to transfer [^3^H]-ADMA across both mouse blood-brain and blood-CSF barriers. Importantly, this is the first study to reveal saturable transport of ADMA at the blood-CSF barrier (choroid plexus).

The decrease in the concentration of [^3^H]-arginine and [^3^H]-ADMA in the presence of unlabelled arginine or ADMA is consistent with inhibition of transporters involved in uptake but it could also be due to stimulation of cellular efflux. As a classical system y^+^ transporter, CAT-1, is bi-directional in its transport activity and can result in the exchange of cationic amino acids between the two sides of the membrane (47)(48). In fact, transport is faster when substrate is present at the opposite (*trans*-) side of the membrane (48). This effect is exhibited by system-y^+^ and called trans-stimulation(49).

Interestingly, in our *in situ* brain perfusion study, transport of [^3^H]-arginine (11.6 nM) across the blood-CNS barriers was insensitive to inhibition by 0.5-100 μM unlabelled ADMA, but sensitive to inhibition by 500 μM unlabelled ADMA. In contrast, the transport of [^3^H]-ADMA (62.5 nM) into all samples was sensitive to inhibition by 100 μM arginine (inhibition ranging from 64.3-80.8%). This is in agreement with studies using human embryonic kidney cells (HEK293) stably overexpressing CAT1 and vector-transfected control cells, which have shown that 100 μM ADMA inhibition of CAT1-mediated transport of *L*-arginine was undetectable (CAT1 having an IC_50_ 758 (460–1251) μM), but 100 μM L-arginine inhibition of CAT1-mediated cellular uptake of ADMA was detectable (CAT1 having an IC_50_ 227 (69–742) μM)(13). This suggests that higher concentrations of unlabelled ADMA (i.e. >100 μM) would be needed to significantly affect [^3^H]-arginine transport by CAT-1 in agreement with our study. In contrast, another study has revealed that 100 μM unlabelled ADMA can exert a significant inhibitory effect on [^3^H]-arginine transport, likely via CAT, into human dermal microvascular endothelial cells (50). Although, as also observed in our study, lower concentrations of ADMA (2.5-10 μM) did not affect [^3^H]-arginine transport in these cells.

Interestingly, cross-competition studies revealed that [^3^H]-arginine uptake into the isolated incubated rat choroid plexus can be inhibited by another methylated arginine, N^G^-methyl-*L*-arginine, at a concentration of 500 μM (51). This *in vitro* method focuses on molecule movement across the apical/CSF side of the choroid plexus, in contrast to the luminal/blood side of the choroid plexus that is examined by the *in situ* brain/choroid plexus perfusion technique.

Interactions between arginine and ADMA have previously been demonstrated in relation to cellular transport (49). There is evidence for both inhibition of uptake and increased efflux of ADMA in the presence of arginine (13,49).

For example, Shin et al. in 2017 demonstrated an enhanced efflux of endogenous arginine and ADMA from a human umbilical vein endothelial cell line due to extracellular exogenous arginine exposure(49). This is consistent with the trans-stimulation of system y^+^ transporters.

Our studies confirm that the transport systems used by ADMA and *L*-arginine at the blood-CNS barriers appear to be shared to some degree, as they each affect the others’ transport, but to a differing extent. Understanding this relationship is of clinical relevance as dietary supplementation with *L*-arginine has been shown to alleviate endothelial dysfunctions caused by impaired NO synthesis (8)(9). Oral supplementation with one dose of 10g arginine has been shown to produce a peak arginine plasma concentration of 200-300 μM (52). This present study would support the theory that arginine supplementation can increase NO production not by inhibiting arginine influx, but by inhibition of [^3^H]-ADMA influx and/ or stimulating the efflux of [^3^H]-ADMA. The reasons for this are as follows. Firstly, our data supports the observations that the transporter for arginine would be operating at nearly maximum capacity in other words is nearly fully saturated (K*_m_* 25-77 μM) within the physiological plasma concentration range of endogenous arginine (human ∼100 μM and mouse ∼140 μM)(29) (28). This together with the facts that: (i) *L-* arginine transport by CAT-1 is a pre-requisite of NO production by the endothelium (53)(54) and ii) eNOS is normally already saturated (K*_m_* ∼3-36 μM) with endogenous intracellular *L*-arginine (840±90 μM)(10)(11)(55) (54), would indicate that raising the plasma concentration of arginine would not increase NO production by increasing the intracellular concentration of arginine (56).

Secondly, our data provides *in situ* evidence that ADMA can cross cell membranes using transporters, transport occurs in both directions and that arginine can reduce ADMA cellular accumulation. Thus, it is possible that supplementation with arginine would affect ADMA transport resulting in the displacement of intracellular ADMA from eNOS (inhibition constant (K*_i_*) 0.9μM) (11) and increased NO production (57,58)(56). ADMA may also increase the apparent K*_m_* of NOS for arginine (57,58). Indeed, arginine supplementation may cause multiple mechanisms to act in parallel ultimately causing an increase in NO production.

## Conclusion

This study has examined the transport of [^3^H]-*L*-arginine and [^3^H]-ADMA across BBB and blood-CSF barrier using physicochemical assessment and an *in situ* mouse brain and choroid plexus perfusion technique. These tripolar cationic amino acids are structurally related but have opposite functions. *L*-Arginine being the exclusive physiological substrate for the NOS family, which synthesizes NO in endothelial cells and neuronal cells, and ADMA being an inhibitor of NOS and so inhibiting NO production.

Results indicate that both [^3^H]-arginine and [^3^H]-ADMA have a gross charge at pH 7.4 of +0.981, but [^3^H]-arginine has a lower lipophilicity than [^3^H]-ADMA. Both [^3^H]-arginine and [^3^H]-ADMA can cross the blood-brain and blood-CSF barriers, but have differing abilities to accumulate in the brain and CSF. This is likely related to their differing ability to use specific transporters for cationic amino acids expressed at these interfaces. This is the first study to present (i) evidence of saturable [^3^H]-ADMA transport at the blood-CSF barrier and (ii) that [^3^H]-ADMA undergoes significant CNS-to-blood efflux across both blood-brain and blood-CSF barriers. Our results also indicate that there is at least some overlap in the specificity of the transporter(s) for arginine and ADMA at the blood-CNS interfaces. For example, unlabelled arginine decreased the CNS accumulation of [^3^H]-ADMA. It is not yet known if this is due to inhibiting [^3^H]-ADMA influx or stimulating [^3^H]-ADMA efflux.

Our data also provides an explanation for the *L*-arginine paradox supporting the hypothesis that increased NO production due to arginine supplementation is likely related to enhanced ADMA efflux rather than increased arginine influx. This is of clinical relevance as NO is an endogenous endothelial vasodilator, and its enhanced production is beneficial in diseases that are associated with endothelial dysfunction.

## DATA SHARING

All data underlying the results are available as part of the article and no additional source data are required.

## ACKNOWLEDGEMENTS

This paper includes data from the PhD thesis of Mehmet Fidanboylu (59). Abstracts of this work have been published (60).

## ABBREVIATIONS

ADMA: asymmetric dimethylarginine or N^G^,N^G^-dimethyl-*L*-arginine
BBB: blood-brain barrier
CAT1: cationic amino acid transporter-1
CNS: central nervous system
CSF: cerebrospinal fluid
CVOs: circumventricular organs
dpm: disintegrations per minute
eNOS: endothelial nitric oxide synthase
K*_m_*: half-saturation constant
V*_i_*: initial volume of distribution
NO: nitric oxide
NOS: nitric oxide synthase
K*_in_*: unidirectional transfer constant

## Supporting information files

**S1 Fig**: **Comparative uptake of [^3^H]-arginine, [^3^H]-ADMA and [^14^C]-sucrose as a function of time measured by *in situ* brain perfusion in anaesthetized mice.** Uptake is expressed as the percentage ratio of tissue to plasma (mL.100 g^-1^). Perfusion fluid contained either [^3^H]-arginine and [^14^C]-sucrose (open markers) or [^3^H]-ADMA and [^14^C]-sucrose (filled markers). Each point represents the mean ± SEM of 4-7 animals (GraphPad Prism 6.0 for Mac).

**S2 Fig: Distribution of [**^3^**H]-arginine, [^3^H]-ADMA and [^14^C]-sucrose in capillary depletion samples as a function of time.** Uptake is expressed as the percentage ratio of tissue to plasma (mL.100 g^-1^). Each point represents the mean ± SEM of 5 animals. Kin and Vi values were determined as the slope and ordinate intercept of the computed regression lines (GraphPad Prism 6.0 for Mac).

**S3 Fig: Comparative distribution of [^3^H]-arginine, [^3^H]-ADMA and [^14^C]-sucrose in the CSF, pineal gland, choroid plexus and pituitary gland following *in* situ brain perfusion as a function of time.** Uptake is expressed as the percentage ratio of tissue or CSF to plasma (mL.100 g^-1^). Each point represents the mean ± SEM of 4-7 animals (GraphPad Prism 6.0 for Mac).

**S4 Fig: The effect of 100μM un-labelled L-arginine on the distribution of [^3^H]-arginine in capillary depletion samples**. Uptake is expressed as the percentage ratio of tissue or CSF to plasma (mL.100 g-1). Brain samples have been corrected for [^14^C]-sucrose (vascular space). Perfusion time is 10 minutes. Each bar represents the mean ± SEM of 6-7 animals. (GraphPad Prism 6.0 for Mac). *p < 0.05, **p < 0.01.

**S5 Fig: The effect of 100μM un-labelled *L*-arginine on the distribution of [^3^H]-arginine in the CSF, choroid plexus and circumventricular organs.** Uptake is expressed as the percentage ratio of tissue or CSF to plasma (mL.100 g-1). Perfusion time is 10 minutes. Each bar represents the mean ± SEM of 6-7 animals (GraphPad Prism 6.0 for Mac). \**p* < 0.05, \*\**p* < 0.01, \*\*\**p* < 0.001.

**S6 Fig: The effect of 100μM un-labelled ADMA on the uptake of [3H]-ADMA in the capillary depletion samples.** Uptake is expressed as the percentage ratio of tissue to plasma (mL.100 g^-1^) and is corrected for [^14^C]-sucrose (vascular space). Perfusion time is 10 minutes. Each bar represents the mean ± SEM of 4 animals (GraphPad Prism 6.0 for Mac). \*\*\**p* < 0.001.

**S7 Fig**: **The effect of 100μM un-labelled ADMA on the uptake of [^3^H]-ADMA in the CSF, choroid plexuses and circumventricular organs.** Uptake is expressed as the percentage ratio of tissue to plasma (mL.100 g^-1^). Perfusion time is 10 minutes. Each bar represents the mean ± SEM of 2-5 animals (GraphPad Prism 6.0 for Mac). *p<0.01, **p<0.05 and \*\*\**p* < 0.001.

**S8 Fig: The effect of unlabelled *L*-arginine on the regional brain uptake of [^3^H]-ADMA (10 minute perfusion).** Uptake is expressed as the percentage ratio of tissue to plasma (mL.100 g^-1^) and is corrected for [^14^C]-sucrose (vascular space). Each bar represents the mean ± SEM of 5 animals (GraphPad Prism 6.0 for Mac).

**S9 Fig: The effect of unlabelled *L*-arginine on the distribution of [^3^H]-ADMA in capillary depletion samples (10 minute perfusion).** Uptake is expressed as the percentage ratio of tissue to plasma (mL.100 g^-1^) and is corrected for [^14^C]-sucrose (vascular space). Each bar represents the mean ± SEM of 5 animals (GraphPad Prism 6.0 for Mac).

**S10 Fig: The effect of unlabelled *L*-arginine on the distribution of [^3^H]-ADMA in CSF, choroid plexus and CVOs (10 minute perfusion).** Uptake is expressed as the percentage ratio of tissue or CSF to plasma (mL.100 g^-1^). Each bar represents the mean ± SEM of 5 animals (GraphPad Prism 6.0 for Mac).

**S11 Fig: Effect of either 100 μM unlabelled ADMA or 100 μM unlabelled *L-* arginine on the respective uptake and distribution of [^3^H]-ADMA in frontal cortex and choroid plexus.** Uptake is expressed as the percentage ratio of tissue or CSF to plasma (mL.100 g-1). Each bar represents the mean ± SEM of 4-5 animals (GraphPad Prism 6.0 for Mac). One-way ANOVA with Dunnett’s post-hoc test comparing means to control ([^3^H]-ADMA only), \*\*\**p* < 0.001).

## References

1. Kobari M, Fukuuchi Y, Tomita M, Tanahashi N, Takeda H. Role of nitric oxide in regulation of cerebral microvascular tone and autoregulation of cerebral blood flow in cats. Brain Res. 1994 Dec;667(2):255–62.

2. Predescu D, Predescu S, Shimizu J, Miyawaki-Shimizu K, Malik AB. Constitutive eNOS-derived nitric oxide is a determinant of endothelial junctional integrity. American Journal of Physiology-Lung Cellular and Molecular Physiology. 2005 Sep;289(3):L371–81.

3. Borroni B, Akkawi N, Martini G, Colciaghi F, Prometti P, Rozzini L, et al. Microvascular damage and platelet abnormalities in early Alzheimer’s disease. J Neurol Sci. 2002 Nov;203–204:189–93.

4. Förstermann U, Closs EI, Pollock JS, Nakane M, Schwarz P, Gath I, et al. Nitric oxide synthase isozymes. Characterization, purification, molecular cloning, and functions. Hypertension. 1994 Jun;23(6_pt_2):1121–31.

5. Vallance P, Leone A, Calver A, Collier J, Moncada S. Accumulation of an endogenous inhibitor of nitric oxide synthesis in chronic renal failure. Lancet. 1992 Mar 7;339(8793):572–5.

6. Vallance P, Leone A, Calver A, Collier J, Moncada S. Endogenous Dimethylarginine as an Inhibitor of Nitric Oxide Synthesis. J Cardiovasc Pharmacol. 1992 Apr;20:S60– 2.

7. Palmer RMJ, Ashton DS, Moncada S. Vascular endothelial cells synthesize nitric oxide from L-arginine. Nature. 1988 Jun;333(6174):664–6.

8. Cooke JP, Andon NA, Girerd XJ, Hirsch AT, Creager MA. Arginine restores cholinergic relaxation of hypercholesterolemic rabbit thoracic aorta. Circulation. 1991 Mar;83(3):1057–62.

9. Clarkson P, Adams MR, Powe AJ, Donald AE, McCredie R, Robinson J, et al. Oral L-arginine improves endothelium-dependent dilation in hypercholesterolemic young adults. Journal of Clinical Investigation. 1996 Apr 15;97(8):1989–94.

10. Baydoun AR, Emery PW, Pearson JD, Mann GE. Substrate-dependent regulation of intracellular amino acid concentrations in cultured bovine aortic endothelial cells. Biochem Biophys Res Commun. 1990 Dec;173(3):940–8.

11. Cardounel AJ, Cui H, Samouilov A, Johnson W, Kearns P, Tsai AL, et al. Evidence for the Pathophysiological Role of Endogenous Methylarginines in Regulation of Endothelial NO Production and Vascular Function. Journal of Biological Chemistry. 2007 Jan;282(2):879–87.

12. Slenzka AUG, HA, and CE. Removal of intracellular asymmetric dimethyl-l-arginine (ADMA) requires system y+L membrane transporter-despite significant activity of the metabolising enzyme dimethylarginine dimethylaminohydrolase (DDAH). Naunyn Schmiedebergs Arch Pharmacol. 2011;383:1–112.

13. Strobel J, Mieth M, Endreß B, Auge D, König J, Fromm MF, et al. Interaction of the cardiovascular risk marker asymmetric dimethylarginine (ADMA) with the human cationic amino acid transporter 1 (CAT1). J Mol Cell Cardiol. 2012 Sep;53(3):392– 400.

14. Watson CP, Pazarentzos E, Fidanboylu M, Padilla B, Brown R, Thomas SA. The transporter and permeability interactions of asymmetric dimethylarginine (ADMA) and L-arginine with the human blood–brain barrier in vitro. Brain Res. 2016;1648.

15. MarvinSketch. MarvinSketch. ChemAxon version 22.9.0. http://chemaxon.com Accessed 2022. 2022;

16. Sanderson L, Khan A, Thomas S. Distribution of suramin, an antitrypanosomal drug, across the blood-brain and blood-cerebrospinal fluid interfaces in wild-type and P-glycoprotein transporter-deficient mice. Antimicrob Agents Chemother. 2007;51(9):3136–46.

17. Triguero D, Buciak J, Pardridge WM. Capillary depletion method for quantification of blood-brain barrier transport of circulating peptides and plasma proteins. J Neurochem. 1990 Jun;54(6):1882–8.

18. Sanderson L, Dogruel M, Rodgers J, De Koning HP, Thomas SA. Pentamidine movement across the murine blood-brain and blood-cerebrospinal fluid barriers: Effect of trypanosome infection, combination therapy, P-glycoprotein, and multidrug resistance-associated protein. Journal of Pharmacology and Experimental Therapeutics. 2009;329(3).

19. Ohno K, Pettigrew KD, Rapoport SI. Lower limits of cerebrovascular permeability to nonelectrolytes in the conscious rat. American Journal of Physiology-Heart and Circulatory Physiology. 1978 Sep 1;235(3):H299–307.

20. Rapoport SI, Ohno K, Pettigrew KD. Drug entry into the brain. Brain Res. 1979 Aug;172(2):354–9.

21. Zloković B V, Begley DJ, Djuricić BM, Mitrovic DM. Measurement of solute transport across the blood-brain barrier in the perfused guinea pig brain: method and application to N-methyl-alpha-aminoisobutyric acid. J Neurochem [Internet]. 1986 May [cited 2016 May 25];46(5):1444–51. Available from: http://www.ncbi.nlm.nih.gov/pubmed/3083044

22. Williams SA, Abbruscato TJ, Hruby VJ, Davis TP. Passage of a delta-opioid receptor selective enkephalin, [D-penicillamine2,5] enkephalin, across the blood-brain and the blood-cerebrospinal fluid barriers. J Neurochem [Internet]. 1996 Mar [cited 2016 May 17];66(3):1289–99. Available from: http://www.ncbi.nlm.nih.gov/pubmed/8769896

23. Gjedde A. High- and Low-Affinity Transport of D-Glucose from Blood to Brain. J Neurochem. 1981 Apr 5;36(4):1463–71.

24. Gjedde A. Calculation of cerebral glucose phosphorylation from brain uptake of glucose analogs in vivo: A re-examination. Brain Res Rev. 1982 Jun;4(2):237–74.

25. Patlak CS, Blasberg RG, Fenstermacher JD. Graphical Evaluation of Blood-to-Brain Transfer Constants from Multiple-Time Uptake Data. Journal of Cerebral Blood Flow & Metabolism. 1983 Mar 29;3(1):1–7.

26. Blasberg RG, Fenstermacher JD, Patlak CS. Transport of α-Aminoisobutyric Acid across Brain Capillary and Cellular Membranes. Journal of Cerebral Blood Flow & Metabolism. 1983 Mar 29;3(1):8–32.

27. Stoll J, Wadhwani KC, Smith QR. Identification of the cationic amino acid transporter (system y+) of the rat blood-brain barrier. J Neurochem. 1993;60(5):1956–9.

28. Böger RH, Bode-Böger SM. The Clinical Pharmacology of L-Arginine. Annu Rev Pharmacol Toxicol. 2001 Apr;41(1):79–99.

29. Hallemeesch MM, Vissers YLJ, Soeters PB, Deutz NEP. Acute reduction of circulating arginine in mice does not compromise whole body NO production. Clinical Nutrition. 2004 Jun;23(3):383–90.

30. Leypoldt F, Choe CU, Gelderblom M, von Leitner EC, Atzler D, Schwedhelm E, et al. Dimethylarginine Dimethylaminohydrolase-1 Transgenic Mice Are Not Protected from Ischemic Stroke. PLoS One. 2009 Oct 7;4(10):e7337.

31. Jung CS, Lange B, Zimmermann M, Seifert V. The CSF concentration of ADMA, but not of ET-1, is correlated with the occurrence and severity of cerebral vasospasm after subarachnoid hemorrhage. Neurosci Lett. 2012 Aug;524(1):20–4.

32. Fickling, Holden, Cartwright, Nussey, Vallance, Whitley. Regulation of macrophage nitric oxide synthesis by endothelial cells: a role for N ^G^, N ^G^-dimethylarginine. Acta Physiol Scand. 1999 Oct 24;167(2):145–50.

33. Kooijmans SAA, Senyschyn D, Mezhiselvam MM, Morizzi J, Charman SA, Weksler B, et al. The Involvement of a Na ^+^ - and Cl ^−^ -Dependent Transporter in the Brain Uptake of Amantadine and Rimantadine. Mol Pharm. 2012 Apr 2;9(4):883–93.

34. del Pino MMS, Peterson DR, Hawkins RA. Neutral Amino Acid Transport Characterization of Isolated Luminal and Abluminal Membranes of the Blood-Brain Barrier. Journal of Biological Chemistry [Internet]. 1995 Jun 23;270(25):14913–8. Available from: http://www.jbc.org/cgi/doi/10.1074/jbc.270.25.14913

35. O’Kane RL, Viña JR, Simpson I, Zaragozá R, Mokashi A, Hawkins RA. Cationic amino acid transport across the blood-brain barrier is mediated exclusively by system y+. Vol. 291, American Journal of Physiology - Endocrinology and Metabolism. Schmitt, U. et al. (2012) ‘In vitro P-glycoprotein efflux inhibition by atypical antipsychotics is in vivo nicely reflected by pharmacodynamic but less by pharmacokinetic changes’, Pharmacology Biochemistry and Behavior, 102(2), pp. 312–320. doi: 10.1016/; 2006. p. E412–9.

36. Smith QR. Transport of Glutamate and Other Amino Acids at the Blood-Brain Barrier. J Nutr. 2000 Apr;130(4):S1016–22.

37. Miller LP, Pardridge WM, Braun LD, Oldendorf WH. Kinetic Constants for Blood— Brain Barrier Amino Acid Transport in Conscious Rats. J Neurochem. 1985 Nov 5;45(5):1427–32.

38. Segal MB, Preston JE, Collis CS, Zlokovic B V. Kinetics and Na independence of amino acid uptake by blood side of perfused sheep choroid plexus. American Journal of Physiology-Renal Physiology. 1990 May 1;258(5):F1288–94.

39. Preston JE, Segal MB. The uptake of anionic and cationic amino acids by the isolated perfused sheep choroid plexus. Brain Res. 1992 May;581(2):351–5.

40. Segal MB, Preston JE, Collis CS, Zlokovic B V. Kinetics and Na independence of amino acid uptake by blood side of perfused sheep choroid plexus. American Journal of Physiology-Renal Physiology. 1990 May 1;258(5):F1288–94.

41. Christensen HN, Antonioli JA. Cationic Amino Acid Transport in the Rabbit Reticulocyte. Journal of Biological Chemistry. 1969 Mar;244(6):1497–504.

42. White MF, Christensen HN. Cationic amino acid transport into cultured animal cells. II. Transport system barely perceptible in ordinary hepatocytes, but active in hepatoma cell lines. Journal of Biological Chemistry. 1982 Apr;257(8):4450–7.

43. White MF, Gazzola GC, Christensen HN. Cationic amino acid transport into cultured animal cells. I. Influx into cultured human fibroblasts. J Biol Chem. 1982 Apr 25;257(8):4443–9.

44. White MF. The transport of cationic amino acids across the plasma membrane of mammalian cells. Biochimica et Biophysica Acta (BBA) - Reviews on Biomembranes. 1985 Dec;822(3–4):355–74.

45. Wang H, Kavanaugh MP, North RA, Kabat D. Cell-surface receptor for ecotropic murine retroviruses is a basic amino-acid transporter. Nature. 1991 Aug;352(6337):729–31.

46. Watson CP, Sekhar GN, Thomas SA. Identification of transport systems involved in eflornithine delivery across the blood-brain barrier. Frontiers in Drug Delivery. 2023 May 23;3.

47. Kim JW, Closs EI, Albritton LM, Cunningham JM. Transport of cationic amino acids by the mouse ecotropic retrovirus receptor. Nature. 1991 Aug;352(6337):725–8.

48. Closs EI, Basha FZ, Habermeier A, Förstermann U. Interference ofL-Arginine Analogues withL-Arginine Transport Mediated by the y+Carrier hCAT-2B. Nitric Oxide. 1997 Feb;1(1):65–73.

49. Shin S, Thapa SK, Fung HL. Cellular interactions between L-arginine and asymmetric dimethylarginine: Transport and metabolism. PLoS One. 2017 May 31;12(5):e0178710.

50. Xiao S, Wagner L, Mahaney J, Baylis C. Uremic levels of urea inhibit <scp>l</scp> - arginine transport in cultured endothelial cells. American Journal of Physiology-Renal Physiology. 2001 Jun 1;280(6):F989–95.

51. Stuhlmiller DFE, Boje KMK. Characterization of <scp>l</scp> -Arginine and Aminoguanidine Uptake into Isolated Rat Choroid Plexus: Differences in Uptake Mechanisms and Inhibition by Nitric Oxide Synthase Inhibitors. J Neurochem. 1995 Jul 23;65(1):68–74.

52. Tangphao O, Grossmann M, Chalon S, Hoffman BB, Blaschke TF. Pharmacokinetics of intravenous and oral L-arginine in normal volunteers. Br J Clin Pharmacol. 1999 Mar 24;47(3):261–6.

53. Zani BG, Bohlen HG. Transport of extracellular <scp>l</scp> -arginine via cationic amino acid transporter is required during in vivo endothelial nitric oxide production. American Journal of Physiology-Heart and Circulatory Physiology. 2005 Oct;289(4):H1381–90.

54. Shin S, Mohan S, Fung HL. Intracellular l-arginine concentration does not determine NO production in endothelial cells: Implications on the “l-arginine paradox.” Biochem Biophys Res Commun. 2011 Nov;414(4):660–3.

55. Hardy TA, May JM. Coordinate regulation of L-arginine uptake and nitric oxide synthase activity in cultured endothelial cells. Free Radic Biol Med. 2002 Jan;32(2):122–31.

56. Tsikas D, Böger RH, Sandmann J, Bode-Böger SM, Frölich JC. Endogenous nitric oxide synthase inhibitors are responsible for the <scp>L</scp> -arginine paradox. FEBS Lett. 2000 Jul 28;478(1–2):1–3.

57. Böger RH. Asymmetric Dimethylarginine, an Endogenous Inhibitor of Nitric Oxide Synthase, Explains the “L-Arginine Paradox” and Acts as a Novel Cardiovascular Risk Factor. J Nutr. 2004 Oct;134(10):2842S–2847S.

58. Bode-Boger S, Scalera F, Ignarro L. The l-arginine paradox: Importance of the l-arginine/asymmetrical dimethylarginine ratio. Pharmacol Ther. 2007 Jun;114(3):295– 306.

59. Fidanboylu M. Blood-CNS transport mechanisms in pathophysiology and drug delivery. [PhD thesis] King’s College London. 2013.

60. Fidanboylu M, Dogruel M, Thomas SA. CAATs at the mouse choroid plexus. Br J Pharmacol [Internet]. 2009 [cited 2023 Dec 7];7(3):036P. Available from: http://www.pa2online.org/abstracts/Vol7Issue3abst036P.pdf

